# Multimodal personalization of transcranial direct current stimulation for modulation of sensorimotor integration

**DOI:** 10.1101/2024.11.06.622280

**Authors:** Jan-Ole Radecke, Alexander Kühn, Tim Erdbrügger, Yvonne Buschermöhle, Sogand Rashidi, Hannah Stöckler, Benjamin Sack, Stefan Borgwardt, Till R. Schneider, Joachim Gross, Carsten H. Wolters, Andreas Sprenger, Rebekka Lencer

**Author notes:** Corresponding author at: Department of Psychiatry and Psychotherapy, University of Lübeck, Ratzeburger Allee 160, 23562, Germany. Declarations of interest: None.

## Abstract

Transcranial direct current stimulation (tDCS) for the modulation of ongoing eye movements provides an ideal model for investigating sensorimotor integration. Within neural networks subserving smooth pursuit eye movements, visual motion area V5 is a core hub to integrate visual motion information with oculomotor control. Here, we applied personalized tDCS explicitly targeting individual V5 in healthy human participants using algorithmic optimization informed by functional magnetic resonance imaging and combined electro- and magnetencephalography. We hypothesized subtle modulation of sensorimotor integration during pursuit and assessed the gain by personalized tDCS targeting V5, compared to personalized tDCS targeting the frontal eye field, as well as conventional normative tDCS over V5. Indeed, pursuit initiation was specifically delayed during personalized cathodal tDCS targeting right V5 indicating the involvement of distinct functional subregions of V5 in the initial sensorimotor integration of visual motion information with pursuit eye movements, but not the maintenance of ongoing pursuit. The results were well-controlled by anodal and sham tDCS, different pursuit tasks, finite-element simulations of individual electric fields, and by two additional control experiments, one that applied personalized tDCS targeting frontal eye field and another that applied normative tDCS over V5. Importantly, in contrast to personalized tDCS targeting FEF and normative tDCS over V5, personalized tDCS targeting V5 effectively modulated pursuit by adapting electric fields to individual anatomical and functional V5 properties. Our results provide evidence for the specific involvement of area V5 in sensorimotor integration during pursuit initiation and the ability of (targeted) tDCS to specifically introduce subtle modulation of the brain network underlying smooth pursuit eye movements. Further, our results indicate the potential of personalized tDCS to alter behavior as the main aspect of interest in human neuromodulation.

## 1. Introduction

Smooth pursuit eye movements (SP) significantly shape our daily interactions with the environment [1] and provide an ideal model for studying sensorimotor integration mechanisms [2,3]. Based on the temporal dynamics of pursuit and the involved brain regions, SP initiation is typically dissociated from sustained SP maintenance. As part of the neural network underlying eye movement control [4,5], the extrastriate visual motion area V5/MT+ [6] has been identified as a key hub for the integration of visual motion processing during both SP initiation and maintenance [2,7–12].

Animal physiology studies demonstrated that the initiation of SP is predicted by activity in the V5-equivalent area MT+ [12–15]. Furthermore, lesion studies of MT+ in macaque monkeys demonstrated impaired SP initiation and maintenance [16–19]. This evidence is complemented by microstimulation of MT+ that increased SP velocities towards the stimulated hemisphere (i.e., ipsiversive SP), and decreased contraversive eye velocities [10]. Accordingly, human lesion studies including area V5 have demonstrated both impaired SP initiation and maintenance [20–22]. In line with this, suprathreshold neuromodulation by transcranial magnetic stimulation (TMS) targeting right V5 in humans induced an overall decrease of ipsiversive (i.e., rightward) SP velocity [23]. Furthermore, functional imaging studies associated activity in the human area V5 with SP stimulus velocity on the one hand as well as the corresponding eye velocity on the other hand, with pronounced V5 activity in the right hemisphere, compared to the left hemisphere [24–26].

Subthreshold neuromodulation of V5 in the healthy brain provides an interesting model for a better understanding of the role of V5 for SP [27]. Furthermore, such techniques can be used to mimic subtle functional impairments of sensorimotor integration, observed in the elderly or clinical populations [28–34]. More specifically, subtle subthreshold neuromodulation by transcranial electric stimulation (tES), including direct current (tDCS) and alternating current stimulation (tACS) can induce transient online-effects and after-effects that outlast the actual stimulation. Several studies induced altered visual motion processing using tDCS or tACS [35–40], but the effect of tDCS on sensorimotor transformation remains elusive. To our knowledge, only one recent study applied normative tDCS over V5 [27] to explicitly modulate sensorimotor integration during SP but did not describe tDCS-induced changes of SP performance.

Conventional tDCS, and tES in general, suffers from a high proportion of non-responders [41–43] and overall small effect sizes in meta-analyses [44,45]. This may partly be explained by the fact that conventional tDCS uses the identical normative scalp electrode montage across subjects. However, based on extensive research on finite element method (FEM) simulations of individual transcranial electric fields [46], it has been shown that, due to volume conduction, inter-individual anatomical differences shape individual tES electric fields that vary near the putative effective threshold to induce subthreshold neural modulation [46–49]. Importantly, neurophysiological markers of tES effects [50–52] and behavior [53–55] have been related to individually simulated electric fields as the putative effector of tES. Based on these challenges of normative tES, it has been suggested to use algorithmic optimization to personalize tES montages, based on explicit assumptions about the individual anatomy and target properties [46,56–58] to control individual transcranial electric fields [59,60]. In the above-mentioned study [27], although no SP modulation by tDCS was described, FEM simulations of transcranial electric fields indicated low directional intensities for the applied normative montage that were significantly smaller compared to personalized electric field intensities. However, actual experimental applications of personalized tES remain sparse [53,61–63], probably due to the resource-intensive approach and, consequently, limited examples for the actual gain of personalized compared to normative tES.

In the present study we applied personalized multichannel tDCS targeting V5 in the right hemisphere (experiment 1) to specifically modulate V5-associated SP initiation and maintenance of ipsiversive (i.e., rightward) SP (Fig. 1). We tested this hypothesis by explicit definition of the individual target V5 and realistic anatomical head models (structural MRI and diffusion weighted imaging, DWI) to compute personalized tDCS montages and thus individually targeted and algorithmically optimized tDCS electric fields. Functional magnetic resonance imaging (MRI) was employed to estimate individual V5 target location and combined electro-(EEG) and magnetoencephalography (MEG; combined MEEG) to estimate target orientation. We assessed the effect of personalized anodal, cathodal and sham tDCS on different aspects of sensorimotor integration during SP, namely SP initiation (as assessed by the step-ramp task, SR), maintenance of continuous SP (TRI task), and top-down control of SP during continuous pursuit with blanking (TRIBL task; Fig. 1 and S3). We hypothesized antagonistic effects of personalized cathodal versus anodal tDCS targeting V5 on ipsiversive SP initiation and maintenance that builds up during tDCS (online-effects) and outlasts the actual stimulation (after-effects). Furthermore, two control experiments were included to test a) the spatial specificity of tDCS targeting V5 compared to personalized tDCS targeting the right frontal eye field (FEF; experiment 2), b) effects of the personalization procedure per se by personalized tDCS targeting remote FEF (experiment 2), and c) the gain by personalized tDCS targeting V5 compared to normative tDCS over V5 (experiment 3). Finally, individual FEM electric fields simulations were analyzed to substantiate experimental findings in an independent analytical way.

**Figure 1.**
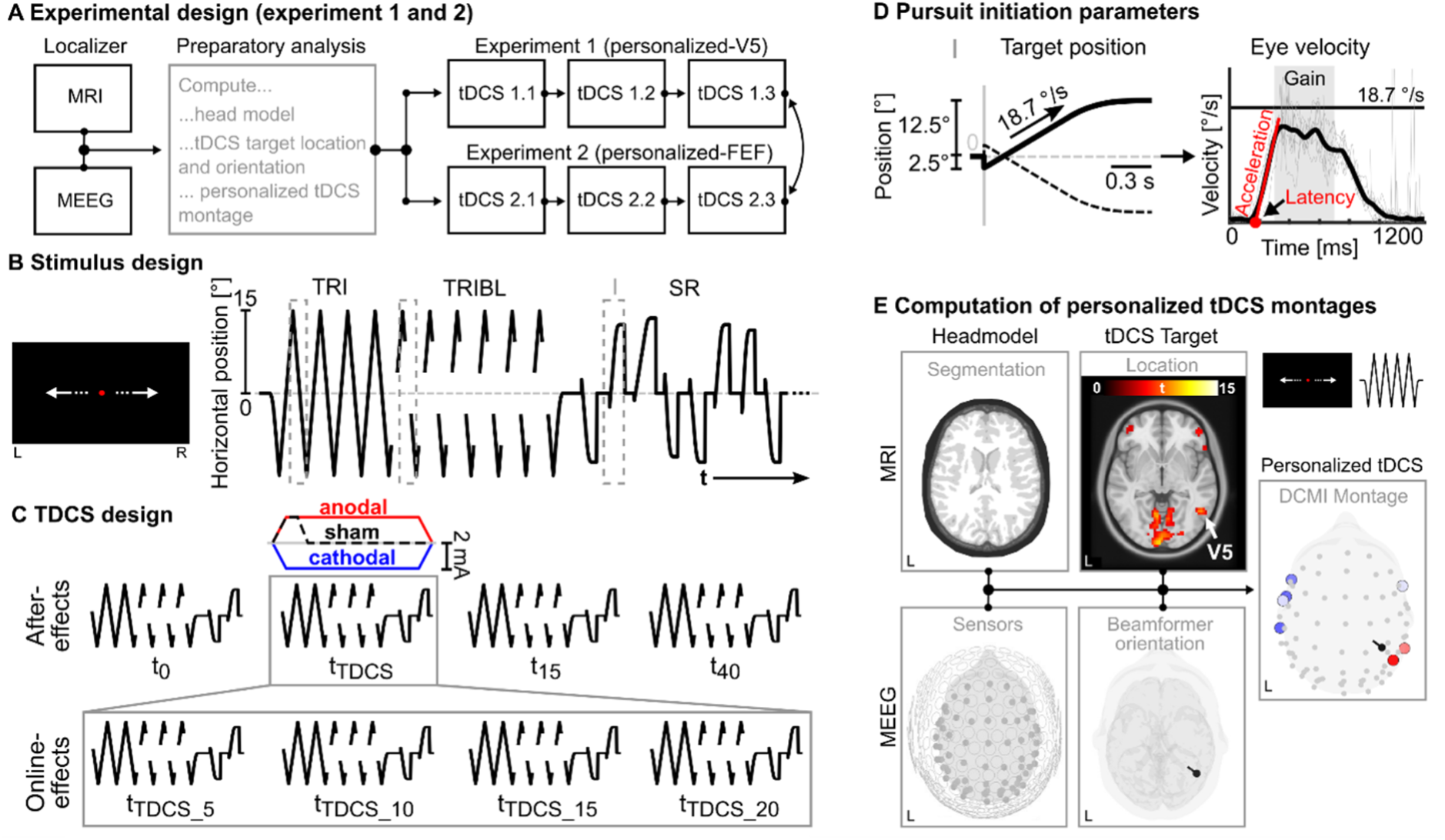
Experimental design. **A)** Personalized tDCS in experiment 1 and 2 comprised eight experimental sessions on separate days per participant (black boxes). During the first two appointments participants completed MRI and MEEG localizer experiments. Localizer data was used to compute personalized tDCS montages (gray box). Sham, anodal, and cathodal tDCS was applied in three subsequent sessions targeting V5 or FEF. After participants completed the first experiment, three further tDCS sessions were completed for the respective second experiment. The initial assignment to experiment 1 or 2, as well as the order of sham, anodal, and cathodal tDCS within each experiment was counterbalanced across participants. **B)** Horizontally moving visual stimuli were presented. To evaluate the effect of personalized tDCS on different aspects of SP, three tasks were presented, namely continuous SP (TRI), continuous SP with blanking (TRIBL), and foveopetal step-ramp stimuli (I, SR). **C)** In each tDCS session, SP tasks were presented before (t_0_), at four timepoints during (t_TDCS_5_, t_TDCS_10_, t_TDCS_15_, t_TDCS_20_), and 15 (t_15_) or 40 minutes (t_40_) after tDCS application. **D)** Pursuit initiation parameters were computed based on the eye tracking data recorded during the step-ramp task. Eye velocities (right) were derived from the eye position (left) as a function of time. Pursuit latency and initial acceleration were computed to define SP initiation performance. Early gain was computed to estimate ongoing SP performance (300 to 700 ms after target motion onset). Target velocity is indicated by a black line at 18.7 °/s. **E)** To compute personalized tDCS montages, structural MRI data were acquired to create a six-compartment segmentation of the individual head (top left). Together with the sensor positions from the combined MEEG session (bottom left), as well as individually calibrated skull conductivity and white matter anisotropy (not shown) individual FEM head models were computed. Individual tDCS targets were defined, using functional MRI data acquired during continuous SP to define the target location in the right hemisphere (top middle, MNI z = -4), and combined MEEG data to define the target orientation (bottom middle). Together the head model and tDCS target information were used to compute personalized tDCS montages with the DCMI algorithm. The workflow to target the area V5 is shown for one exemplary subject.

We aimed to assess the effects of novel multimodally personalized tDCS targeting area V5 for the subtle neuromodulation of SP behavior. Indeed, in contrast to personalized tDCS targeting FEF and normative tDCS over V5, cathodal personalized tDCS targeting right V5 specifically modulated sensorimotor integration during SP initiation, but not SP maintenance during ongoing eye movements. The multimodal personalization and analysis of tDCS electric fields ensured the control of observed inter-individual anatomical and functional variability. Further, results from three experiments allowed conclusions about the distinct involvement of V5 in the modulation of SP initiation, specifically by personalized tDCS targeting this area.

## 2. Results

### 2.1 Personalized tDCS adapts to inter-individual target variability

Individual stimulation targets showed varying location and orientation (Fig. 2A-B). Specific SP-related BOLD activity was observed in individually estimated V5 locations near previously reported V5 coordinates (x/y/z MNI-coordinates, V5: 45/-67/2 ± 3/6/4; Tab. S1) [7,64]. Individual V5 activation maxima deviated by 7 ± 3 mm to the mean location, yielding a considerable inter-individual variability of tDCS target locations (Fig. 2B). Second-level analysis of BOLD activity revealed a significant increase of V5 activity during continuous pursuit, compared to fixation in a cluster in inferior and middle occipital, as well as middle and inferior temporal cortex of the right hemisphere (cluster size *k* = 97, cluster *p* = .007, local maximum: 42/-73/2, *t*_18_ = 7.6) [65]. For illustration, individual statistical maps from the Pursuit versus Fixation contrast were interpolated on the MNI brain, and FDR-corrected one-sample *t*-tests are shown (Fig. 2A). A mask of the V5-equivalent cytoarchitectonic area hOc5 was additionally applied and confirmed a significantly increased BOLD activity in V5 during SP compared to fixation (*t*_18_ = 6.3, *p* < .001, center coordinates: 45/-61/2) [6]. In line with the heterogenous cortical structural organization of V5, the estimated stimulation target orientations showed a variety of quasi-radial, and quasi-tangential orientations across participants (Fig. 2B).

**Figure 2.**
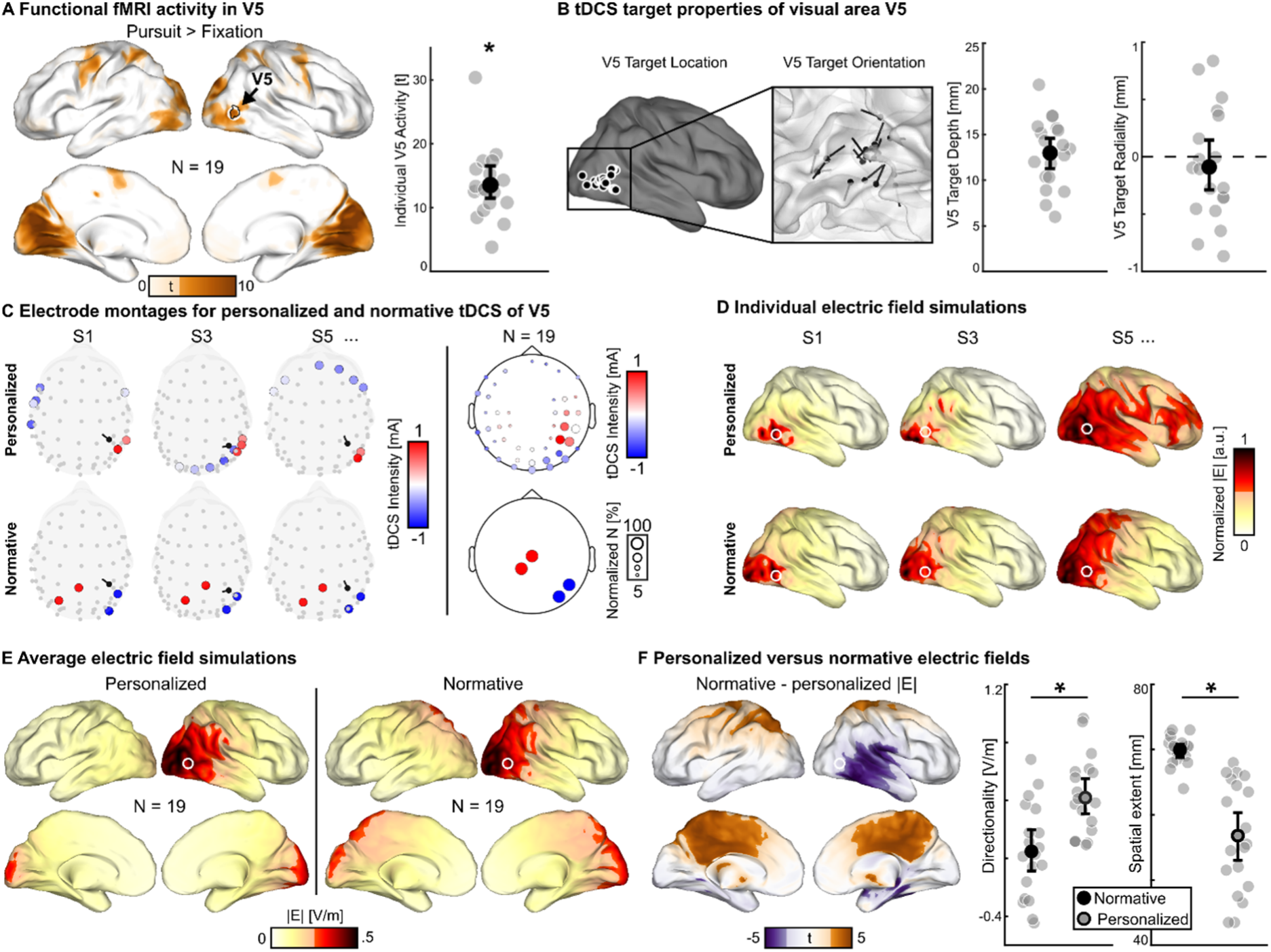
Target V5 and personalized transcranial electric fields. **A)** Left: Second-level BOLD activity during ongoing SP, compared to fixation, revealed a significant cluster that corresponds to visual area V5. The V5-equivalent cytoarchitectonic area hOc5 (black lines) corresponds to the average coordinate of individual V5 estimates (white circle). FDR-corrected *t*-values are shown, interpolated on the inflated cortical surface of the MNI brain. Right: Individual target V5 activity. Bootstrap mean and confidence interval is depicted together with individual local maxima *t*-values of the first-level statistical contrast of BOLD activity during ongoing SP, compared to fixation. **B)** Target V5 locations (left) are shown on an inflated cortical surface of the MNI brain. Estimated V5 orientations (middle) are shown together with the white matter surface of the MNI brain. Right: V5 target depth relative to the inner skull surface and radiality (1/-1 = radial/anti-radial, 0 = tangential) bootstrap means and confidence intervals are depicted together with individual values. **C)** Personalized tDCS montages targeting individual V5 (top) and normative tDCS montages over V5 (bottom) are shown for three exemplary subjects (left; top view). Topographical representations for the whole sample are shown for personalized (top right) and normative tDCS montages (bottom right). Circle sizes illustrate the normalized number of individual montages that comprised the respective electrode, color illustrates the average tDCS intensity per electrode across the whole sample. **D)** Electric field simulations for the same three subjects and personalized (top) and normative (bottom) tDCS montages, respectively. Normalized electric field intensities |**E**| were interpolated on the inflated right cortical surface of the MNI brain, with values below 50% of the maximal intensity masked (light colors). Individual tDCS target locations in the right V5 are indicated by white circles. **E)** Lateral and medial view of average simulated electric fields for personalized (left) and normative (right) tDCS interpolated on the inflated cortical surface of the MNI brain, with values below 50% of the maximal intensity masked (light colors). The average tDCS target location in the right V5 is indicated by a white circle, respectively. **F)** Left: Comparison between personalized and normative simulated electric fields revealed increased electric field intensities in bilateral parietal and medial brain regions for normative tDCS, whereas personalized electric field intensities were increased in right temporal, and parietal regions anterior to the average V5 location (white circle), corresponding to more anterior individual V5 locations (compare panel B). Lateral and medial view of significant *t*-values are shown (bold colors), with *t*-values corresponding to FDR-corrected *p*-values > 0.05 masked (light colors). Right: Directionalities in the target V5 for electric fields induced by normative tDCS were significantly smaller and spatial extent of the electric field across the cortex was significantly larger, compared to personalized target directionalities and spatial extent. Bootstrap means and confidence intervals are depicted together with individual values. * indicate *p* < .05.

Personalized tDCS montages targeting individual right V5 confirmed a clear adaptation to the target location and orientation, placing cathodes on the scalp roughly over the target in the direction of estimated target orientation. Across subjects, exemplary for the application of personalized cathodal tDCS, a cluster of anodes was observed including electrode positions P6, P8, and CP6, whereas a cluster of cathodes was observed including electrode positions PO8, PO10, O2, and O10. However distinct target orientations and quasi-radial target orientations are reflected by varying tDCS electrode positions across the scalp (Fig. 2C) [60]. Overall SP performance was high in the expected range for young healthy participants (Tab. 2, see Supplement) [27].

**Table 1.** Sample information for the personalized and the normative tDCS participants. For both samples sample size (N), age (M ± SD), gender (frequencies), handedness (frequencies), raw values for the Beck Depression Inventory (BDI-II 126) and percentile ranks for the Multiple Choice Vocabulary Test (MWT-B 127) are provided. F = female, M = male, R = right-handed, L = left-handed, A = ambidextrous.

| Parameter | Personalized tDCS<br>targeting V5 and FEF | Normative tDCS<br>over V5 |
| --- | --- | --- |
| Sample size | 19 | 19 |
| Age | 26.2 ± 7.6 | 25.5 ± 7.1 |
| Gender | F: 11, M: 8 | F: 11, M: 8 |
| Handedness | R: 16, L: 2, A: 1 | R: 16, L: 3 |
| BDI-II | 1.7 ± 2.2 | 1.4 ± 1.2 |
| MWT-B | 47.7 ± 19.6 | 55.2 ± 20 |
| Days between tDCS sessions | 7.9 ± 7.8 | 8.5 ± 4.7 |

**Table 2.** Descriptive smooth pursuit performance. Great grand average pursuit performance for experiment 1 (V5-personalized), experiment 2 (FEF-personalized), and experiment 3 (V5-normative). Mean (± SD) is reported for the respective parameters of continuous pursuit (TRI), continuous pursuit with blanking (TRIBL), and foveopetal step-ramps (SR). a.u. = arbitrary unit.

| Task | Parameter | Unit | Experiment 1<br>(V5-personalized) | Experiment 2<br>(FEF-personalized) | Experiment 3<br>(V5-normative) |
| --- | --- | --- | --- | --- | --- |
| **TRI** | Maintenance gain | a.u. | 0.957 ± 0.042 | 0.936 ± 0.044 | 0.941 ± 0.039 |
| **TRIBL** | Preblank gain | a.u. | 0.839 ± 0.108 | 0.819 ± 0.108 | 0.814 ± 0.112 |
| | Deceleration latency | ms | 130 ± 17 | 134 ± 22 | 134 ± 19 |
| | Deceleration | $^{\circ}/s^2$ | 47 ± 9 | 46 ± 9 | 54 ± 7 |
| | Residual gain | a.u. | 0.439 ± 0.223 | 0.425 ± 0.216 | 0.314 ± 0.145 |
| | Re-acceleration latency | ms | -132 ± 35 | -136 ± 39 | -136 ± 39 |
| | Re-acceleration | $^{\circ}/s^2$ | 27 ± 10 | 26 ± 10 | 26 ± 9 |
| | Postblank gain | a.u. | 0.811 ± 0.125 | 0.786 ± 0.106 | 0.731 ± 0.145 |
| **SR** | Pursuit latency | ms | 160 ± 13 | 163 ± 13 | 154 ± 11 |
| | Acceleration | $^{\circ}/s^2$ | 128 ± 34 | 126 ± 33 | 109 ± 29 |
| | Early maintenance gain | a.u. | 0.827 ± 0.086 | 0.809 ± 0.087 | 0.799 ± 0.094 |

### 2.2 Personalized tDCS targeting V5 specifically modulates pursuit initiation

Personalized tDCS was applied to modulate SP performance depending on tDCS condition (sham, anodal, cathodal), timepoint (t_0_, t_TDCS_5_, t_TDCS_10_, t_TDCS_15_, t_TDCS_20_, t_15_, t_40_) and SP stimulus direction (leftward, rightward). Personalized tDCS targeting V5 specifically modulated SP latencies in the SR task, as revealed by an omnibus analysis (three-way interaction: *p* = .044; Fig. 3A-C, Tab. S2). Additional LMM analyses indicated an online effect during tDCS (*p* = .015; Fig. 3A-C, Tab. 3), but no tDCS after-effects (*p* = .484; Tab. 4). Specifically personalized cathodal tDCS delayed rightward SP latencies for t_TDCS_20_, relative to t_TDCS_5_ (*p* = .002, *d*_z_ = 0.93). A follow-up analysis confirmed that overall SP latencies across all timepoints of the sham session (t_0_, t_TDCS_5_ to t_TDCS_20_, t_15_, t_40_) were delayed for leftwards compared to rightwards SP (*p* = .007, *d*_z_ = 0.64; compare Fig. 3A), indicating that personalized cathodal tDCS targeting the right V5 hampered latencies of rightward SP to the level of initially delayed leftward SP latencies. In addition, a paradoxical effect by personalized cathodal tDCS was observed during t_TDCS_15_ between leftward and rightward SP latencies (*p* = .002, *d*_z_ = 0.89) and latencies for leftward SP during t_TDCS_15_ were significantly delayed for personalized cathodal tDCS, compared to sham (*p* = .042, *d*_z_ = .72; Fig. 3A).

**Figure 3.**
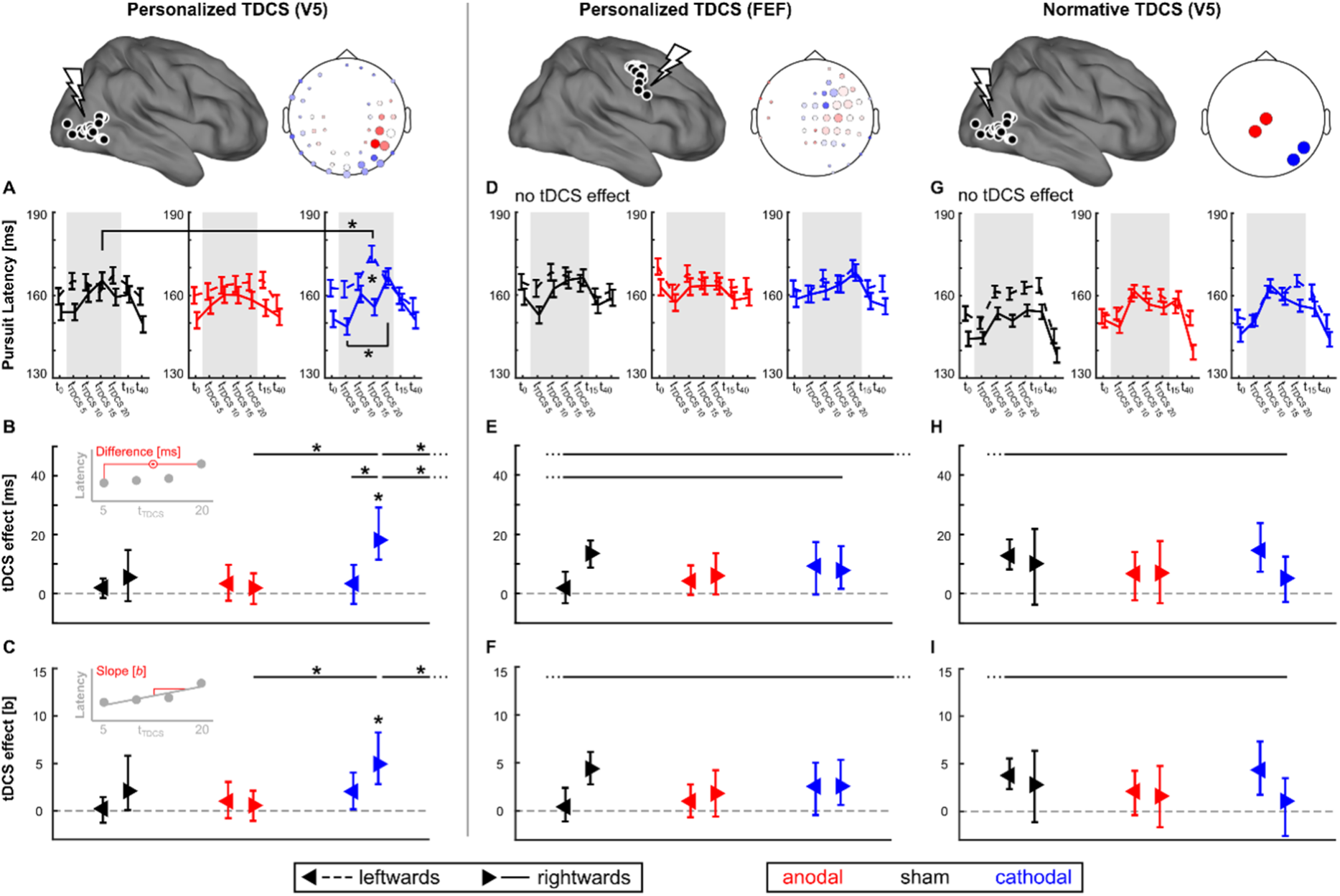
tDCS effects on SP latencies during the step-ramp task. **A) to C)** In experiment 1 (V5-personalized), significant modulation of SP latencies during the SR task by personalized cathodal tDCS targeting V5 was observed. An effect was observed across **A)** all timepoints (t_0_, t_TDCS_5_, t_TDCS_10_, t_TDCS_15_, t_TDCS_20_, t_15_, t_40_). Gray shaded boxes indicate the 20 minutes tDCS interval. LMM analyses revealed an online effect of personalized cathodal tDCS with significantly delayed latencies for ipsiversive (i.e., rightwards) SP during the last timepoint (t_TDCS_20_), compared to the first (t_TDCS_5_) timepoint during t_TDCS_. **B)** tDCS effect estimated as difference between t_TDCS_20_ versus t_TDCS_5_ is plotted for sham, anodal, and cathodal tDCS conditions, as well as leftwards and rightwards SP latencies, respectively (bootstrap mean ± confidence interval). Top left inset illustrates the computation of the difference tDCS effect. Follow-up analysis of difference tDCS effects reveals significant differences for personalized cathodal tDCS for rightward SP latencies, compared to leftward SP latencies of the same tDCS condition, rightward SP latencies during anodal tDCS, as well as an increased difference tDCS effect of personalized cathodal tDCS targeting V5, compared to personalized tDCS targeting FEF and normative cathodal tDCS over V5. **C)** tDCS effect estimated as slope of robust regression across all t_TDCS_ timepoints is plotted for sham, anodal, and cathodal tDCS conditions, as well as leftwards and rightwards SP latencies, respectively (bootstrap mean ± confidence interval). Top left inset illustrates the computation of the slope tDCS effect. Slope estimates of tDCS effects are significantly increased for personalized cathodal tDCS targeting V5, compared to anodal tDCS, as well as normative cathodal tDCS over V5. **D) to F)** No specific tDCS effects were observed for personalized tDCS targeting the right FEF. **G) to I)** No specific tDCS effects were observed for normative tDCS over the right V5. * indicate *p* < .05 (Bonferroni-corrected). tDCS targets and topographical illustration of the cathodal tDCS montage are depicted for the experiment 1 (V5-personalized, top left), experiment 2 (FEF-personalized, top middle), and experiment 3 (V5-normative, top right), respectively.

**Table 3.** Personalized tDCS targeting V5 induces online effects modulating SP initiation. Results of LMM analysis indicate significant transient modulation of SP initiation, specifically pursuit latency and initial acceleration during SR, by personalized tDCS targeting V5 during the tDCS application (online effects, i.e. considering timepoints t_TDCS_5_, t_TDCS_10_, t_TDCS_15_, and t_TDCS_20_). TRI = continuous pursuit, TRIBL = continuous pursuit with blanking, SR = step-ramp. * indicate *p*-values < .05.

| Task | Parameter | tDCS condition |  | tDCS condition * Timepoint |  | tDCS condition * Direction |  | tDCS condition * Timepoint * Direction |  |
| --- | --- | --- | --- | --- | --- | --- | --- | --- | --- |
| | | F | $p$ | F | $p$ | F | $p$ | F | $p$ |
| TRI | Maintenance gain | 1.35 | .263 | 0.26 | .955 | 1.71 | .183 | 0.63 | .705 |
| TRIBL | Preblank gain | **6.09 \*** | **.003** | 1.31 | .252 | 0.55 | .58 | 0.28 | .948 |
|  | Deceleration latency | 1.59 | .207 | 0.19 | .98 | 1.37 | .257 | 0.28 | .945 |
|  | Deceleration | 2.61 | .077 | 0.15 | .989 | 0.12 | .886 | 0.86 | .521 |
|  | Residual gain | 0.32 | .73 | 0.26 | .955 | 0.54 | .586 | 0.11 | .995 |
|  | Re-acceleration latency | **4.79 \*** | **.01** | 0.7 | .648 | 0.77 | .467 | 1.1 | .36 |
|  | Re-acceleration | 0.02 | .978 | 0.28 | .945 | 0.39 | .675 | 0.59 | .736 |
|  | Postblank gain | **3.24 \*** | **.041** | 0.39 | .887 | 0.02 | .982 | 0.46 | .836 |
| SR | Pursuit latency | 0.63 | .534 | 1.11 | .355 | 0.79 | .456 | **2.7 \*** | **.015** |
|  | Initial acceleration | 0.29 | .751 | 0.77 | .596 | 0.25 | .78 | **2.56 \*** | **.02** |
|  | Early maintenance gain | **4.74 \*** | **.01** | 0.41 | .87 | 0.15 | .862 | 0.74 | .62 |

**Table 4.** No tDCS after-effects. Results of LMM analysis do not indicate a significant modulation of SP parameters after personalized tDCS targeting V5 (after effects, i.e. considering timepoints t_0_, t_15_, and t_40_). TRI = continuous pursuit, TRIBL = continuous pursuit with blanking, SR = step-ramp. * indicate *p*-values < .05.

| Task | Parameter | tDCS condition |  | tDCS condition * Timepoint |  | tDCS condition * Direction |  | tDCS condition * Timepoint * Direction |  |
| --- | --- | --- | --- | --- | --- | --- | --- | --- | --- |
| | | F | $p$ | F | $p$ | F | $p$ | F | $p$ |
| TRI | Maintenance gain | 0.71 | .491 | 1.56 | .159 | 0.05 | .954 | 0.19 | .979 |
| TRIBL | Preblank gain | 0.45 | .638 | 1.19 | .312 | 1 | .369 | 0.46 | .838 |
|  | Deceleration latency | 0.87 | .42 | 0.58 | .748 | 0.06 | .943 | 0.9 | .492 |
|  | Deceleration | 0.18 | .836 | 0.37 | .896 | 0.92 | .4 | 0.31 | .934 |
|  | Residual gain | 0.51 | .599 | 0.27 | .95 | 0.1 | .902 | 0.15 | .99 |
|  | Re-acceleration latency | **6.7 \*** | **.002** | 1.94 | .074 | 0.08 | .924 | 0.54 | .782 |
|  | Re-acceleration | 0.44 | .644 | 0.11 | .995 | 0.62 | .542 | 1.42 | .207 |
|  | Postblank gain | 1.34 | .264 | 1.14 | .342 | 0.04 | .963 | 0.15 | .989 |
| SR | Pursuit latency | 0.72 | .487 | 0.53 | .786 | 0.07 | .936 | 0.92 | .484 |
|  | Initial acceleration | 0.28 | .755 | 0.48 | .824 | 0.01 | .989 | 0.43 | .857 |
|  | Early maintenance gain | 1.36 | .259 | 0.83 | .545 | 0.24 | .784 | 0.14 | .991 |

To follow-up tDCS online effects for rightward SP latencies (Fig. 3A), these were defined both as difference between the first and last timepoint during t_TDCS_ (t_TDCS_20_ – t_TDCS_5_; 18 [12, 29] ms, bootstrapped mean and [95%-confidence interval]; Fig. 3B), as well as the slope of a linear fit across all four t_TDCS_ timepoints (t_TDCS_5_ to t_TDCS_20_; 4.9 [2.8, 8.3] ms/5 minutes ; Fig. 3C). While the difference tDCS effect can be directly related to the significant post-hoc contrasts of the LMM analysis, the slope tDCS effect yields a more robust estimate considering the paradoxical effect but thereby introduces potential noise from the SP latencies observed during t_TDCS_10_ and t_TDCS_15_. Therefore, we consider the difference and slope estimates of the tDCS effect as complementary. Significant differences between personalized anodal and cathodal tDCS for the rightward SP latencies were revealed for both difference (*z* = -3, *p* = .008, *d*_z_ = 0.72) and slope estimates (*z* = -2.58, *p* = .03, *d*_z_ = 0.61) of tDCS effects. In addition, difference tDCS effects differed between leftwards and rightwards SP latencies during personalized cathodal tDCS (*z* = -2.41, *p* = .024, *d*_z_ = 0.59). The corresponding effect for the slope estimate did not survive Bonferroni-correction (*p* = .11). In addition, the difference tDCS effect was significantly larger for personalized cathodal tDCS targeting V5, when compared to personalized tDCS targeting FEF (*z* = 2.08, *p* = .019, *d*_z_ = 0.5) while the same effect for the slope estimate missed significance (*p* = .057).

During tDCS, an online effect on initial acceleration was also observed (*p* = .02; Tab. 3), however, this effect was not indicated by the preceding omnibus analysis and post-hoc tests did not survive correction for multiple comparisons.

Besides the modulation of SP latencies, no specific tDCS effects (i.e., tDCS condition * timepoint or tDCS condition * timepoint * direction interaction effects) were observed for tasks indicating sustained SP maintenance performance with continuously visible targets (TRI), top-down mechanisms involved in ongoing SP with temporary target blanking (TRIBL) or ongoing SP performance during the SR task following SP initiation (all *p* > .523; Tab. 3-4, Tab. S2). Furthermore, neither personalized tDCS targeting the right FEF as a control region (within-group, *p* = .862; Fig. 3D-F, Tab. S4), nor normative tDCS over V5 (*p* = .93; between-group, matched sample of N = 19; Fig. 3G-I, Tab. S5) induced a modulation of SP latencies during tDCS comparable to the observed effects for personalized tDCS targeting V5 (Fig. 3A-C). Finally, the results were not compromised by individual tDCS side-effects (see Supplement).

### 2.3 Personalized tDCS targeting V5 increases tDCS efficacy compared to normative tDCS over V5

To evaluate the electric field differences presumably leading to distinct tDCS effects, electric field simulations were compared between the applied personalized tDCS targeting V5 and the normative V5 montage for the N = 19 subjects from experiment 1 (personalized V5; Fig. 2C-F). Electric field simulations showed comparable target intensities (*p* = .209, normative |E|_IT_ = 0.62 ± 0.38 V/m, personalized |E|_IT_ = 0.66 ± 0.45 V/m, M ± SD). However, normative electric fields across the cortex showed significantly increased intensities in non-target dorsal and medial parts of bilateral parietal cortex compared to personalized electric fields. In contrast, personalized electric fields showed increased intensities in the right lateral occipital-parietal-temporal cortex compared to the normative electric fields (Fig. 2F). In addition, directionalities were significantly increased for personalized compared to normative electric fields (*z* = -4.87, *p* < .001, *d*_z_ = 1.11; normative |E|_DIR_ = 0.05 ± 0.32 V/m, personalized |E|_DIR_ = 0.42 ± 0.28 V/m, M ± SD; Fig. 2F). Electric field spatial extent was significantly decreased for personalized, compared to normative tDCS (*z* = 6.81, *p* < .001, *d*_z_ = 1.74 normative |E|_EXTENT_ = 70 ± 2 mm, personalized |E|_EXTENT_ = 57 ± 8 mm, M ± SD; Fig. 2F). In conclusion, personalized electric field topographies were shifted towards individual tDCS targets in the right V5, thus effectively considering the varying individual tDCS target locations and orientations in the right V5 and reducing co-stimulation of non-target parietal cortex, compared to the normative electric fields.

Indeed, the evaluation of tDCS effects on ipsiversive SP latencies revealed significantly increased tDCS effects for personalized cathodal tDCS targeting V5 compared to normative tDCS over V5 (difference tDCS effect: *z* = 2.1, *p* = .018, *d*_s_ = 0.7; slope tDCS effect: *z* = 1.81, *p* = .036, *d*_s_= 0.61).

## 3. Discussion

We conducted a comprehensively controlled experiment to assess the effects of personalized tDCS targeting V5 on SP initiation and maintenance as a direct behavioral read-out of tDCS effects on sensorimotor integration. We found a spatially and functionally specific, significant modulation of SP initiation by personalized cathodal tDCS that was absent during anodal or sham tDCS. This effect was independent from possible practice effects [27] and tDCS side-effects (Supplement). Furthermore, electric field simulations and a control experiment applying normative tDCS over V5 signify more precise electric fields and an increased gain by personalized tDCS in contrast to normative tDCS.

### 3.1 Personalized tDCS targeting V5 specifically modulates pursuit initiation

To interpret our key finding of specific SP initiation modulation (i.e., SP latencies) by personalized cathodal tDCS, it is important to consider the multimodal role of the human visual area V5/MT+. V5 can be functionally dissociated into the middle temporal visual area (MT) and the medial superior temporal area (MST) [7,9]. In the macaque monkey, it has been shown that MT receives bottom-up visual information [66,67] and integrates these to signal visual motion on the retina as a velocity signal during SP [11]. MST receives projections from MT [67] and MST itself maintains reciprocal connections to various visual areas, as well as parietal and premotor nodes of the oculomotor network [66,68] that govern top-down signals during ongoing SP, e.g., predicting target motion even in the absence of a visual target [11]. Importantly, neural activity in MT+ is involved in SP initiation, but relies on gain control by higher-order network nodes such as the FEF [12,13,15]. In line with these studies, human V5/MT+, comprising putative MT and MST, has been suggested as a key hub for the sensorimotor transformation during SP [7,69], thus integrating bottom-up visual motion information with top-down oculomotor control during SP initiation and SP maintenance.

In the present study, SP initiation (SP latency in the SR task) was hampered by personalized cathodal tDCS targeting V5. The observed impairment of ipsiversive, but not contraversive, SP is in line with previous TMS modulation of the right V5 in humans [23]. Animal studies described an impairment of ipsiversive SP for foveopetal step-ramps after lesions of the foveal representation of MT [17,19] anduman lesion studies including V5 described similar ipsiversive SP deficits [20–22]. However, the same authors also differentiate contralesional SP deficits (retinotopic) from directional deficits that affect SP ipsiversive to the lesion site [17,20–22]. During the step of the applied SR task, the visual target is moved 2.5 ° to one side, before it starts to move in the opposite direction with a constant velocity ramp. This paradigm allows subjects to start pursuit without an initial saccade as soon as the visual target crosses the point of the subject’s central fixation at the start of each trial. However, this also implies that in the time between the step onset and the midline-crossing of the visual target, subjects fixate the center of the screen and thus are presented a retinotopic visual stimulus moving from the step eccentricity towards the participant’s central fixation, before the visual target is then fixated during the actual pursuit. Thus, in the present experiment personalized cathodal tDCS presumably induced retinotopic deficits that affected sensorimotor transformation after the leftward target step onset moved the visual target to the hemifield contralateral to the targeted right V5, but before the onset of rightward SP. This would explain the observed isolated deterioration of SP latencies that might follow retinotopic deficits in the hemifield contralateral to the targeted right V5. At the same time initial acceleration (SR task), SP maintenance (TRI task) and top-down aspects of SP in the blanking task (TRIBL task), which would rather indicate directional effects [18], remained unaffected [21]. Retinotopic SP deficits could indicate the involvement of peripheral representations in MT, in contrast to foveal MT representations, and/or adjacent MST parts. Lesions of the latter also result in directional SP deficits [16,17,19]. In contrast to tDCS targeting V5, no SP modulation was observed by tDCS targeting right FEF, further marking the spatial specificity of personalized tDCS targeting right V5. We conclude that in our experiment personalized cathodal tDCS specifically modulated early sensorimotor transformation within V5 putative subregion MT, or feedforward communication from MT to MST. We further speculate, either that sensorimotor transformation of foveated visual motion information was not affected by personalized cathodal tDCS, or that tDCS-induced initial directional disturbances were compensated by higher-order cognitive mechanisms coming into play under maintenance of ongoing SP [28,29,70–72].

### 3.2 Personalized cathodal tDCS targeting V5 transiently reduces cortical excitability

Conventionally, cathodal tDCS has been defined as placing the cathode (anode) over the assumed target region to reduce (increase) cortical excitability [73]. Among other mechanistic explanations for tDCS [74], it has been proposed that tDCS electric fields induce cortical membrane polarizations that lead to a subthreshold modulation of neuronal firing-rates and altered cortical excitability [73,75], in addition to changes of blood flow [76] and local GABA/glutamate metabolism at the stimulation site [77–79]. Cathodal tDCS specifically was described to reduce cortical excitability [73,75] and local glutamate concentration [78]. Therefore, personalized cathodal tDCS in the present study presumably reduced cortical excitability in V5, thereby hampering the integration of visual motion information to elicit SP initiation as indicated by delayed SP latencies.

We observed a transient modulation of SP latencies by single-bout personalized cathodal tDCS (online-effect). No after-effects were observed 15 minutes after tDCS or thereafter. Instead, the online tDCS effect on SP initiation evolved slowly over the 20 minutes tDCS resulting in a significant latency difference between the first (t_TDCS_5_) and the last (t_TDCS_20_) read-out timepoints. Immediate tDCS modulations were previously shown to induce NMDA-receptor mediated long-term plasticity (LTP) or long-term depression (LTD) that drive longer-lasting tDCS effects [80–82]. Thus, we conclude that personalized cathodal tDCS in the present study induced immediate changes in MT excitability which exerted a statistically robust tDCS effect on behavioral SP performance only when tDCS-induced LTD-like effects were established after several minutes of continuous tDCS.

In previous studies, both anodal and cathodal tDCS over V5 induced an improvement in the discrimination accuracy of coherent motion perception in visual scenes [40]. Following previous speculations [37], Battaglini and colleagues [40] suggested that, on the one hand, anodal tDCS increases coherent motion discrimination by increasing the excitability of neurons in the putative subregion MT of V5 that are tuned to correlated motion signals. On the other hand, cathodal tDCS reduces the overall excitability of MT, thereby relatively reducing the response to uncorrelated noise in competing motion scenes with low coherence. At the same time, cathodal tDCS also reduces the discrimination of coherent motion signals in visual scenes with high signal-to-noise ratio. In line with this evidence, we observed a specific impairment of the direct and naturalistic SP sensorimotor response to a single moving visual target without competing motion noise (i.e., with high signal-to-noise ratio) during personalized cathodal tDCS. We further speculate that we did not observe performance facilitation by personalized anodal tDCS in the present young, healthy participants due to a ceiling effect [83] of their anyway high SP performance, in addition to practice-associated improvements obtained during the study (Supplement) [27]. In contrast, impaired SP latencies observed in the elderly and clinical populations [28–30], might be improved by anodal tDCS representing interesting target populations to further examine the clinical benefits of (personalized) tDCS for future studies.

Taken together, by applying personalized, i.e., targeted tDCS and differentiating distinct functional properties of SP, we can pinpoint the modulation of SP latencies by personalized cathodal tDCS to an early phase of sensorimotor transformation, presumably involving reduced excitability and LTD-like modulation of V5 subregion MT that delays SP performance.

### 3.3 Normative tDCS over V5 does not modulate pursuit initiation

In contrast to the specific modulation of ipsiversive SP latencies during personalized tDCS targeting V5, no online effect was observed by normative tDCS over V5, extending previous findings [30] that normative tDCS over V5 did not result in any tDCS after-effects on SP [27]. However, simulations of tDCS electric fields in a small sample (N = 6) already suggested rather small target directionalities by normative tDCS that could be increased by personalized tDCS montages with respect to individual targets. Here, we confirm, in a much larger sample, that normative tDCS over V5 yields limited target directionalities, compared to personalized tDCS targeting V5. This improvement of simulated personalized compared to normative electric fields is in line with previous findings of increased personalized directionalities when targeting the parietal cortex [60], the somatosensory cortex [61] and increased electric fields across the brain and within the target region for personalized electric fields targeting the auditory cortex [63]. Furthermore, we extend previous findings by whole-brain analysis of personalized compared to normative simulated electric fields indicating reduced spatial extent of the targeted electric fields and a shift of the electric field topography away from bilateral non-target parietal regions and towards individual target V5 estimates in temporal regions within the right hemisphere (Fig. 2E-F).

Although it has been proposed repeatedly to implement electric field simulations and optimization in tES protocols [48,56–58,84], actual applications of personalized tES remain rare [53,61–63]. Thus, little is known about the gain of personalized, compared to conventional normative tES. Therefore, in addition to the analysis of simulated electric fields, we investigated whether the observed behavioral online effect of personalized cathodal tDCS on ipsiversive SP latencies were specific to the applied personalization approach, by re-analyzing data from a matched sample where normative tDCS was applied over the right V5. In contrast to personalized tDCS targeting V5, we did not observe a modulation of SP during normative tDCS over V5. A more detailed analysis indicated a significantly increased tDCS effect on SP by personalized cathodal tDCS, compared to normative cathodal tDCS. We thus follow that the increased theoretical efficacy of personalized compared to normative tDCS of V5 as indicated by electric field simulations holds true for the actual modulation of SP latencies as direct behavioral read-out of the tDCS-effect. Only a few previous studies provided data from applied personalized tDCS [61] or tACS [53,62,63] and only two of those provided a direct comparison of personalized versus normative tES [61,63]. To our knowledge, as of now, no study described the gain by personalized compared to normative tDCS with respect to the modulation of behavior. Khan and colleagues [61] found a significant modulation of somatosensory evoked activity by personalized tDCS targeting the primary somatosensory cortex in ten subjects, but no effect by normative tDCS. In the same study, personalized transcranial electric fields optimized with the DCMI algorithm [85] reached simulated target directionalities of 0.38 ± 0.21 V/m, compared to 0.26 ± 0.13 V/m for normative tDCS (2 mA). In line with these results, our present data indicate that personalized tDCS using the DCMI algorithm targeting V5 shifts the electric field topography towards the explicitly defined tDCS target and thereby increases target directionalities above the effective threshold at the group-level. Normative tDCS may not produce reliable results, due to the relatively limited and untargeted electric fields. Especially in the light of a growing interest in tDCS for clinical application [86,87], our present findings support the huge potential of personalized tES approaches directing individual transcranial electric fields to the assumed cortical target, thereby maximizing the individual stimulation efficacy.

In another study, the application of 2 Hz tACS targeting the auditory cortex (stimulation intensity: ±0.5 mA) induced a phase-specific modulation of frequency-modulated sounds by personalized tACS, but also for normative tACS [63]. In this light, it must be noted that personalized tES does not necessarily increase effectiveness, compared to normative tES. Indeed, normative tES can greatly reduce the resources required for experimental and clinical applications considering additional data acquisition and analysis for target localization, individual head models, and electrode positions needed for the optimization and careful application of personalized tES [48]. To conclude, we suggest that personalized and normative approaches should be handled complementary: By specific assumptions about the individual anatomy and the individual functional tES targets, personalized tES is predestined to test hypotheses about the functional mechanisms underlying specific tES applications, for example the role of V5 in the modulation of SP, as shown in the present study. Naturally, personalized tES combines the analysis of tES read-out parameters with simulated electric fields and neurophysiological data that can inform and extend the analysis. These insights can then be used to validate implicit assumptions of normative tES applications or to *a priori* inform the optimization of next-generation, resource-efficient and effective normative tES montages, as a complementary approach to data-intensive *a posteriori* optimization of normative tES montages [55,88].

In this study, our main finding shows that personalized cathodal tDCS targeting right V5 specifically delayed latencies of ipsiversive SP due to a transient but LTD-like modulation of early sensorimotor transformation in the MT subregion of V5. Personalized tDCS targeting the right FEF as a control region did not modulate pursuit performance indicating the spatial specificity of personalized tDCS targeting V5 and suggest that these effects cannot be explained by the personalization procedure per se. Further, FEM simulations of electric fields suggest that the observed gain of (effective) personalized tDCS, compared to (ineffective) normative tDCS of V5 might be attributed to topographical shifts of personalized transcranial electric fields towards the targeted right V5 at group-level, as well as increased target directionalities and reduced spatial extent. We suggest that personalized tDCS, and tES in general, might be an interesting methodological perspective to increase tES efficacy, especially in clinical research and therapeutical settings.

## 4. Materials and methods

### 4.1 Participants and procedure

Twenty healthy participants were recruited for the application of personalized tDCS targeting V5 (experiment 1) and FEF (experiment 2), respectively. One participant did not complete the experiment due to personal reasons. Thus, complete datasets of nineteen participants were included in the analyses (Tab. 1). To compare results from personalized tDCS to the application of normative tDCS, a subsample from a previous study applying normative tDCS over V5 [27] was statistically matched to the personalized tDCS sample with respect to age, gender, laterality index, MWT-B percentile ranks and BDI-II raw values (paired *t*-tests, all *p* ≥ .25; Tab. 1) and data was re-analyzed. All participants reported no history of psychiatric or neurological disorders and no psychotic disorders of first-degree family members. Participants had normal or corrected-to-normal visual acuity (1 ± 0.1) and gave written informed consent in line with the declaration of Helsinki prior to the experiment. The study was approved by the ethics committees of the Universities of Lübeck and Münster (#20-459 and #2015-263-f-S).

In a full within-subject design, participants completed a battery of SP tasks during eight experimental sessions (Fig. 1; single-blind, see Supplement). During the first session, structural MRI data was recorded to compute individual FEM head models. Additionally, fMRI activity was recorded while subjects [89,90] performed a SP task to localize areas V5 and FEF, in each participant individually. During the second session, combined MEEG was recorded while SP was performed to define the target neuronal orientation at the fMRI-based locations of V5 and FEF [89]. In addition, MEEG data was recorded during electrical stimulation of the median nerve to estimate individual skull conductivities based on somatosensory evoked MEEG activity [90] to inform combined MEEG source analysis and target orientation determination. In the following sections, we describe how we used the resulting individually skull-conductivity calibrated FEM head models (section 4.2) to compute personalized tDCS montages for the individual targets in V5 and FEF (section 4.3 and 4.4). Subsequently, sham-controlled personalized anodal and cathodal tDCS targeting areas V5 and FEF was applied in six subsequent experimental sessions (section 4.5; tDCS conditions and target area were pseudo-randomly assigned) to modulate SP performance (section 4.6). In addition, personalized tDCS targeting V5 was compared to normative tDCS over V5, with respect to simulated transcranial electric fields and SP performance under modulation by tDCS (section 4.7). In section 4.8 the statistical analysis is described.

### 4.2 Calibrated individual FEM head models

Structural MRI data were recorded using a 3-T Siemens Magnetom Skyra scanner (Siemens, Germany) and a 64-channel head coil. In addition to T1- and T2-weighted images (T1: 3D MP-RAGE sequence, TR = 2300 ms, TE =3.6 ms, TI = 1100 ms, FA= 8°; T2: spin echo pulse sequence, TR = 3200 ms; TE = 408 ms, FA = 120°; both at 1 x 1 x 1 mm resolution and 192 x 256 x 256 mm FoV), DWI data were acquired for 64 direction vectors (turbo spin echo sequence, 69 volumes, 1 x 1 x 1 mm resolution, 100 x 100 x 72 mm FoV, SMS with slice acceleration factor 4, TR = 5500 ms, TE = 128 ms, FA = 90°, b = 1000 s/mm^2^) and five additional volumes were acquired with diffusion sensitivity b = 0 s/mm^2^.

Individual geometry-adapted six-compartment hexahedral FEM head models were computed, including calibrated skull conductivities and white matter conductivity anisotropy [27,91] with the DUNEuro toolbox (see Supplement for details). Conductivities were set to normative values for scalp (0.43 S/m), CSF (1.79 S/m), gray matter (0.33 S/m) and white matter (0.14 S/m). For the EEG lead field, skull conductivities were individually calibrated (skull compacta: 0.013 ± 0.01 S/m, skull spongiosa: 0.0468 ± 0.036 S/m; see Supplement) [90,91]. Finally, the EEG lead fields for estimation of the tDCS target orientations, and individual geometry matrices for the tDCS optimization were computed using the calibrated skull conductivities.

### 4.3 Functional localization of V5 and FEF tDCS targets

Personalized tDCS montages were computed based on individual estimates of the stimulation targets in areas V5 (experiment 1) and FEF (experiment 2) of the right hemisphere, respectively (Fig. 1E, Fig. 2) [27], as described in more detail below.

First, to determine the target locations of V5 and FEF using fMRI, blood oxygen level dependent (BOLD) activity was recorded (simultaneous multislice (SMS) with acceleration factor 4, 307 Volumes, TR = 980 ms, TE = 30 ms, FA = 70°, resolution 3 x 3 x 3 mm, 68 x 68 x 56 mm) while participants performed SP (red dot as SP target, diameter 0.5°, black background; 1920 x 1080 screen resolution, 60 Hz refresh rate, NordicNeuroLab, Norway) and eye movements were recorded (Eyelink 1000Plus, 1000 Hz sampling rate; SR Research Ltd., Canada). Participants completed a test battery of SP-tasks similar to the tasks during the tDCS experiments (see Supplement). Each pursuit block was preceded by fixation intervals with a centrally presented red dot (12 s, diameter 0.5°). Functional images were slice-timing corrected, smoothed (6 mm FWHM Gaussian kernel), and co-registered to the normalized T1 image. Individual locations of visual area V5 and FEF in the right hemisphere were determined as local maxima of BOLD activity during SP near putative V5 and FEF regions (Tab. S1) [7,64] in contrast to fixation intervals (*t*-test, *p* < .05 across 5 adjacent voxels, FWE-corrected).

Second, individual target orientations were estimated using beamforming at the right V5 and FEF target locations defined by functional MRI. Therefore, combined MEEG data were recorded (EEG: 60 channels, Easycap, Germany; MEG: 275 axial gradiometers, VSM MedTech Ltd., Canada; 600 Hz sampling rate) while participants completed a continuous pursuit task (TRI; 18.7°/s, ±15° amplitude, 20 blocks of each 8 ramps to the left and right interleaved with short breaks). Before the first block and after the 10^th^ block, participants fixated a red dot in the center of the screen for 15 s. Data were filtered (0.1 Hz highpass-filter, 50 Hz notch filter considering harmonics), cut into 320 epochs (0 to 1.59 s length relative to ramp onsets to left and right), and demeaned. Invalid epochs were rejected semi-automatically (rejected trials: 15.3 ± 11.3 %) and the EEG data was re-referenced to the average reference. tDCS target orientations were estimated using linearly constrained minimum variance (LCMV) beamforming with Unit-Noise-Gain constraint (regularization parameter λ = 5%), following the procedure described in [89] (see Supplement for details).

Electrode positions for the tDCS montage optimization were recorded (FASTRAK, Polhemus Inc., VT/USA). These 74 electrode positions follow the 10-20-system, embedded in a standard EEG cap layout (Easycap, Germany), of which 60 electrodes were used as EEG sensors.

### 4.4 Computation of personalized tDCS montages and electric field simulations

Based on individual anatomical information, integrated in the head models and individual target information from the localizer tasks, personalized tDCS montages were computed to target the area V5 (experiment 1) or FEF (experiment 2), respectively. To this end, the distributed constrained maximum intensity (DCMI) algorithm [53,60,61,85] was employed to compute the personalized tDCS montage that maximizes the injected current at the target location along the target orientation (i.e., the directionality; see Supplement for details). Resulting montages described personalized cathodal tDCS montages. Herein, we define the applied personalized tDCS montages relative to the estimated target orientations, i.e., personalized cathodal tDCS results in montages with the cathodes placed on the scalp roughly in the direction of the estimated target orientation. For personalized anodal tDCS, the polarity of each electrode was inverted. This definition does not resemble normative tDCS conventions where the cathodes/anodes are placed on the scalp directly over the proposed target location defining cathodal/anodal tDCS. This convention implicitly assumes quasi-radial targets but does not consider tangential targets or inter-individual target localization variability. The target orientations in the present study were estimated as the MEEG summed dipole orientation that might not directly reflect the local cortical structure, but rather functional properties of the target orientation. Therefore, the definition of personalized cathodal/anodal tDCS considers individual target locations and orientations.

### 4.5 Application of personalized tDCS

During tDCS (t_TDCS_) personalized anodal, cathodal, or sham tDCS was applied targeting the right V5 and FEF in six pseudo-randomized sessions, separated by at least four days (7.9 ± 7.8 days, M ± SD; Fig. 1C). During each session, 20 minutes of tDCS was applied using a Starstim 8 device (Neuroelectrics, Spain). Small Ag/AgCl electrodes (NG Pistim, 3.14 cm^2^ surface) were placed in custom empty EEG caps (EasyCap, Germany) with the same layout and cap size as used for the EEG measurement and electrode registration to ensure an accurate placement of the stimulation electrodes according to the optimized personalized montages (Fig. 1E). During personalized anodal and cathodal tDCS, a total of 2 mA current was applied (10 s ramp-on and ramp-off). Sham tDCS was applied by the same montages as used for anodal stimulation, but tDCS was applied for only 30 s in addition to the current ramps.

To minimize transcutaneous side-effects, anesthetic creme (2.5 % lidocaine, 2.5 % prilocaine, Anesderm®, Pierre Fabre, Germany) was used [92]. After each t_TDCS_ interval, participants completed a questionnaire asking for their subjectively perceived transcutaneous side-effects during t_TDCS_ (itching, warmth, stitching, throbbing, and pain) on a five-point scale (0 = absent, weak, moderate, pronounced, 4 = intense) [93]. Furthermore, participants rated subjective fatigue for four timepoints during t_TDCS_ on a five-point scale (1 = very alert, awake, moderate, tired, 5 = very tired).

### 4.6 Eye tracking and analysis of eye movement data

During the six tDCS sessions, participants were seated 65 cm in front of an LCD monitor in a closed room with lights off and completed three SP tasks to assess different aspects of SP initiation and maintenance (see Supplement for details) [27]. In short, eye movements were recorded using a video-based eye tracker (Eyelink 1000Plus, SR Research Ltd., Canada). During all tasks, participants were instructed to follow a horizontally moving SP target on the monitor as precisely as possible (red dot with 0.5° diameter, black background; Fig. 1B). During the first task, eight horizontal constant velocity ramps were presented as continuous triangular waveform stimulus (TRI, 18.7°/s, ± 15° amplitude; Fig. 1B-C, Supplement). In the second task, 12 similar horizontal velocity ramps were presented with target blanking between 300 to 1000 ms relative to ramp onset (18.7°/s, ± 15° amplitude, triangular waveform stimulus with blanking, TRIBL; Fig. 1B-C, Supplement). To specifically assess SP initiation and early SP maintenance eight horizontal foveopetal step-ramp stimuli (SR) were presented [94], pseudo-randomly directed either to the left or the right side of the screen. Step-ramps started with a 2.5° step which was immediately followed by a velocity ramp in the opposite direction (18.7°/s, ± 15° amplitude; Fig. 1B-D). To minimize expectation effects, two of eight step-ramps were presented with alternating velocities (9.7 °/s, ± 1.3 ° step-size; 26.7 °/s, ± 3.5 ° step-size). The three tasks were presented in mixed blocks [33] including three task repetitions during each block. This resulted in 24 continuous ramps (TRI), 36 ramps with blanking (TRIBL) and 24 step-ramps (SR) per block.

Effects of tDCS on SP were assessed by presenting each one block of SP tasks before (t_0_), and 15 and 40 minutes after (t_15_, t_40_; after-effects) tDCS application in each of the six experimental sessions. Online effects of tDCS were assessed by presenting four repetitions of SP task blocks equally distributed across the 20 minutes tDCS application (t_TDCS_; i.e., within the first 5, 10, 15, or 20 minutes of tDCS, t_TDCS_5_, t_TDCS_10_, t_TDCS_15_, t_TDCS_20_; Fig. 1C). During t_TDCS_, SP task blocks were separated by simple oculomotor tasks to activate the oculomotor system during the stimulation, while providing an active rest for the participants at the same time. Overall, this resulted in 504 continuous ramps (TRI), 756 ramps with blanking (TRIBL) and 504 step-ramps (SR) that were analyzed for each subject during experiment 1 (V5) and experiment 2 (FEF), respectively.

SP data analysis focused on SP velocity, computed as the first derivative of eye position. Continuous data were epoched (-100 to 1600 ms for TRI and TRIBL and -100 to 1300 ms for SR relative to the ramp onset) and invalid epochs were rejected (experiment 1 (V5): 1.5 ± 2.1 % epochs were rejected; see Supplement). Median eye velocity across epochs was computed separately for each subject, tDCS condition (anodal, cathodal, sham), timepoint (t_0_, t_TDCS_5_, t_TDCS_10_, t_TDCS_15_, t_TDCS_20_, t_15_, t_40_) and stimulus direction (leftward, rightward). SP velocity parameters were computed to assess different aspects of SP performance during SP initiation (SR: SP latency, initial eye acceleration; Fig. 1D) as well as ongoing SP (SR: early maintenance gain; TRI: maintenance gain, gain = eye velocity/target velocity; TRIBL: preblank, residual and postblank velocity gains, deceleration and deceleration latency at the start of the blanking interval, as well as re-acceleration and re-acceleration latency at the end of the blanking interval) [27,71].

### 4.7 Comparing personalized tDCS targeting V5 and normative tDCS over V5

Results from personalized tDCS targeting V5 (experiment 1) were compared to results from the application of normative tDCS over V5 (experiment 3; Tab. 1). The normative application of V5 was designed identically to the personalized procedure with respect to the SP test battery and the overall application of tDCS [27] and data from experiment 3 was re-analyzed according to the analysis procedure in experiment 1. However, instead of personalized tDCS montages, normative tDCS montages were used in experiment 3 that did not vary across subjects, thus also no MEEG and MRI data were acquired before the normative tDCS experiment. In short, to apply normative tDCS, two stimulation electrodes were placed over area V5 of the right hemisphere (PO8, P8) with two central return electrodes (Cp1, Cz) [36,37] according to the 10-20 system. During normative tDCS, 2 mA current was applied for 20 minutes, restricted to 1 mA per electrode. Eye movements were recorded during the same timepoints as in the personalized experiment (t_0_, t_TDCS_5_, t_TDCS_10_, t_TDCS_15_, t_TDCS_20_, t_15_, t_40_). Since SP during t_TDCS_ were not analyzed in detail by Radecke et al. [27], the data from the matched sample were re-analyzed for all tDCS timepoints (t_TDCS_5_, t_TDCS_10_, t_TDCS_15_, t_TDCS_20_) to compare online effects on the SP performance by normative versus personalized tDCS.

Furthermore, electric field simulations were compared between normative and personalized montages to estimate the differences in the potential efficacy between the two tDCS methods, given the individual estimates of the V5 tDCS target. Therefore, normative electric field simulations were computed for the N = 19 subjects from the personalized tDCS sample, using the normative electrode placement, since data for these participants for computation of individual head models and estimation of individual tDCS targets were available. To quantify individual transcranial electric field parameters (see Supplement for details), for each subject, we computed the target intensity (|E|_IT_) the target intensity corrected for the parallelity between the stimulation target orientation vector and the target electric field orientation vector (directionality, |E|_DIR_) [53,60,61], as well as the spatial extent of the electric field relative to the stimulation target (|E|_EXTENT_) [53,60]. The overall cortical electric field (|**E**|) and electric field parameters with respect to the individual stimulation targets (|E|_IT_, |E|_DIR_, |E|_EXTENT_) were compared between personalized versus normative tDCS. For illustration, individual electric fields were interpolated on a common MNI cortical grid and averaged across subjects (Fig. 2).

### 4.8 Statistical analysis of tDCS effects

To analyze tDCS effects on SP performance, linear mixed models (LMMs) were fitted to the data (IBM SPSS Statistics, IBM Corp., USA) for each estimated SP parameter and separately for experiment 1 (V5-personalized), experiment 2 (FEF-personalized), and experiment 3 (V5-normative). In a full within-subject design, tDCS condition (anodal, cathodal, sham), timepoint (t_0_, t_TDCS_5_, t_TDCS_10_, t_TDCS_15_, t_TDCS_20_, t_15_, t_40_) and the visual stimulus direction (leftward, rightward) were included as fixed effect factors. To control inter-individual variability of SP performance, subject ID was included as random effects factor (random intercept; identity covariance structure). tDCS condition, timepoint and ramp direction were considered as repeated factors (first-order autoregressive covariance structure). Main and interaction effects were analyzed in a saturated model. In case of specific tDCS effects (i.e., an interaction between tDCS condition, timepoint, and ramp direction), separate LMM analyses were computed for timepoints of online effects (t_TDCS_5_, t_TDCS_10_, t_TDCS_15_, t_TDCS_20_) and (after-)effects relative to t_0_ (t_0_, t_TDCS_, t_15_, t_40_, using the averaged performance across all t_TDCS_ timepoints), further including tDCS condition and ramp direction as factors.

The significance level was set to *α* = .05 and post-hoc contrasts of estimated marginal means were computed for the highest-order interaction or main effect (Bonferroni-corrected for multiple comparisons). *F*-values and *p*-values are reported for significant main or interaction effects. For post-hoc tests, Bonferroni-corrected *p*-values, mean and standard error of the mean are reported. Effect sizes were estimated as the absolute value of Cohen’s 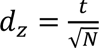 with *N* being the respective sample size and *t* being the *t*-value computed based on the mean difference and standard error of the mean of the difference between estimated marginal means of the respective paired post-hoc contrast [95].

Furthermore, significant LMM tDCS condition * timepoint * stimulus direction interactions indicate specific tDCS effects. To follow-up these results in more detail, tDCS online effects were defined as the difference between the first and the last timepoint during t_TDCS_ (t_TDCS_20_ - t_TDCS_5_) of the respective SP parameter, irrespective of the temporal evolution of this effect. A second, complementary estimate of tDCS effects was computed as the slope (*b*) of a robust linear regression computed across all timepoints during t_TDCS_ (t_TDCS_5_ to t_TDCS_20_) assuming a quasi-linear evolution of the tDCS effect during the stimulation (Fig. 3). Both tDCS effect measures were compared between stimulus directions for each tDCS condition, as well as between tDCS conditions for each stimulus direction using bootstrapped paired *t*-tests (*n* = 10000 bootstrap samples were computed to determine the H_0_ distribution). In addition, tDCS effects for the respective tDCS condition and stimulus direction were compared across experiments using bootstrapped paired *t*-tests to compare between experiment 1 (V5-personalized) versus experiment 2 (FEF-personalized) and unpaired *t*-tests to compare experiment 1 (V5-personalized) versus experiment 3 (V5-normative). For all bootstrap *t*-tests *z*-values and *p*-values (Bonferroni-corrected) are reported and Cohens *d*_z_ (paired contrast) or *d*_s_ (unpaired contrast) was computed based on the observed values [95].

Electric field simulations were compared between normative versus personalized tDCS montages. Individually simulated electric fields were interpolated on an MNI cortical surface representation and paired *t*-tests were computed for every position of the surface grid between personalized and normative tDCS (FDR-corrected) [96]. Furthermore, separate bootstrap paired *t*-test were computed (one-sided) with 10000 iterations using MATLAB (The Mathworks Ltd., MA/USA) for |E|_IT_, |E|_DIR_, and |E|_EXTENT_. For each test, *z*-value, *p*-value and descriptive means and standard errors of the means are reported. Effect size *d*_z_ was estimated based on the observed values [95].

## Abbreviations

SP: Smooth pursuit eye movements
tES: transcranial electric stimulation
tACS: transcranial alternating current stimulation
tDCS: transcranial direct current stimulation
FEM: finite element method
DWI: diffusion weighted imaging
MRI: magnetic resonance imaging
EEG: electroencephalography
MEG: magnetoencephalography
MEEG: combined EEG and MEG
FEF: frontal eye field
BOLD: blood oxygen level dependent
SMS: simultaneous multislice
TRI: continuous pursuit task
LCMV: linearly-constrained minimum variance
DCMI: distributed constrained maximum intensity
TRIBL: continuous pursuit task with blanking
SR: foveopetal step-ramp task
LMM: linear mixed model
MT: middle temporal visual area
MST: medial superior temporal area;

## CRediT authorship contribution statement

**Jan-Ole Radecke:** Conceptualization, Methodology, Software, Investigation, Formal analysis, Writing - original draft, Visualization. **Alexander Kühn:** Investigation, Data curation, Formal Analysis, Writing – review & editing. **Tim Erdbrügger and Yvonne Buschermöhle:** Investigation, Methodology, Software, Writing - review & editing. **Sogand Rashidi and Hannah Stöckler:** Investigation, Data curation, Writing – review & editing. **Benjamin Sack:** Investigation, Writing - review & editing. **Stefan Borgwardt:** Resources, Supervision, Writing - review & editing. **Till R. Schneider:** Supervision, Methodology, Writing - review & editing. **Joachim Gross:** Supervision, Funding acquisition, Writing - review & editing. **Carsten H. Wolters:** Supervision, Project administration, Funding acquisition, Writing - review & editing. **Andreas Sprenger:** Methodology, Software, Supervision, Writing – review & editing. **Rebekka Lencer:** Project administration, Funding acquisition, Conceptualization, Writing - review & editing.

## Acknowledgements

This work was supported by the German Research Foundation (DFG; WO1425/10-1 to CHW, GR2024/8-1 to JG and LE1122/7-1 to RL) and by the Bundesministerium für Gesundheit (BMG; ZMI1-2521FSB006, under the frame of ERA PerMed as project ERAPERMED2020-227). JG was further supported by the DFG (GR2024/11-1, GR2024/12-1, TRR393). JOR was funded by the Clinician Scientist program of the Section of Medicine of the University of Lübeck. We thank Dr. Martin Göttlich and Andreas Wollbrink for technical support, Karin Wilken, Hildegard Deitermann, Ute Trompeter, Susanne Schellbach and Mandy-Josephine Reichhardt for support with the data acquisition, and Prof. Peter Trillenberg for helpful discussions.

