## Supplementary Materials for "Multimodal personalization of transcranial direct current stimulation for modulation of sensorimotor integration"

### S1 Computation of calibrated individual FEM head models

Individual geometry-adapted six-compartment hexahedral FEM head models were computed, including calibrated skull conductivities and white matter conductivity anisotropy [1,2]. T1 and T2 images were co-registered using FSL FLIRT [3]. SPM12[1] was used to segment gray matter, white matter (CAT12) [4], and scalp from the T1 image, as well as skull compacta and cerebrospinal fluid (CSF) from the T2 image [5]. Otsu thresholding [6] was performed on the eroded skull compacta compartment to isolate an additional skull spongiosa compartment from the T2-weighted hypointense skull compacta. Custom MATLAB (The Mathworks Ltd., MA/USA) scripts were applied to further process the six compartment masks using Boolean and morphological operations [2,7,8]. The volume was cut 4 cm below the skull using an axial plane [9]. Next, geometry-adapted hexahedral meshes were computed (0.33 node shift) [10]. After removing eddy current and nonlinear susceptibility artifacts from DWI data using FSL and HySCO [11,12], anisotropic conductivity tensors were computed for the white matter compartment based on an effective medium approach [13]. Lead fields were computed using the Venant source modeling approach implemented in the DUNEuro toolbox [2], based on an equally spaced 2 mm source space grid including only gray matter grid points that meet the Venant condition [14,15].

### S2 Estimation of skull conductivities to compute calibrated head models

For the EEG lead fields, skull conductivities were individually calibrated [2,16]. Therefore, combined MEEG data were recorded (EEG: 60 channels, Easycap, Herrsching, Germany; MEG: 275 axial gradiometers, VSM MedTech Ltd., Vancouver, Canada; 600 Hz sampling rate) during electrical wrist stimulation of the median nerve (1932 monophasic electrical square-wave pulses of length 0.5 ms; inter-stimulus interval was uniformly jittered between 350 and 450 ms; stimulus strength was adjusted relative to the individual threshold such that the right thumb moved clearly) to evoke somatosensory evoked activity. After preprocessing (Filter: 20 to 250 Hz, 50 Hz notch filter considering harmonics; Epochs from -50 to 150 ms relative to onset of the electrical pulses; Trials semi-automatically rejected: 10 ± 5.3 %) and averaging of the data across trials, individual P20/N20 components were determined [17]. An MEG dipole scan was performed, and the resulting dipole location was fixed (residual variance 0.058 ± 0.025). We then minimized the residual variance for the EEG dipole orientation with fixed location [18] by only changing the skull conductivity (residual variance 0.135 ± 0.066). The resulting conductivities for skull compacta ranged from 0.003 to 0.032 S/m (0.013 ± 0.01 S/m)

[1] https://www.fil.ion.ucl.ac.uk/spm

[2] https://www.medizin.uni-muenster.de/duneuro/startseite.html

[2]. The ratio between skull compacta and skull spongiosa was set to 3.6 resulting in conductivities of 0.0468 ± 0.036 S/m for skull spongiosa.

#### S3 Functional localization of V5 and FEF targets

To define tDCS targets, brain activity was recorded to estimate individual V5 and FEF locations (see main manuscript). SP tasks during functional MRI recording included continuous pursuit (TRI, 18.7°/s, ±15° amplitude, four blocks of each eight leftwards and rightwards

**Table S1. Target locations for V5 and FEF.** Location vectors were determined based on the contrast computing larger BOLD activity during continuous pursuit compared to central fixation. In few cases, the contrast between all pursuit tasks, compared to central fixation was used (italic). MNI-coordinates [x, y, z] and T-values of local maxima are depicted for the right V5 and the right FEF during continuous pursuit for all subjects and averaged across subjects (* indicate *p* < .05 across at least 5 adjacent voxels, FWE-corrected).

| ID | V5 | | | | FEF | | | |
|---|---|---|---|---|---|---|---|---|
| | x | y | z | T | x | y | z | T |
| **S1** | 48 | -58 | -4 | 10.8* | 45 | -1 | 50 | 10.8* |
| **S2** | _42_ | _-64_ | _5_ | _3.8*_ | 57 | 5 | 41 | 9.2* |
| **S3** | 48 | -70 | -1 | 13.5* | 48 | -4 | 50 | 9.9* |
| **S4** | 42 | -79 | -1 | 7.4* | _51_ | _5_ | _32_ | _7.5*_ |
| **S5** | 48 | -64 | 2 | 12.4* | _57_ | _11_ | _32_ | _8.3*_ |
| **S6** | 42 | -70 | -4 | 8.4* | _42_ | _-1_ | _56_ | _11.5*_ |
| **S7** | 42 | -67 | 2 | 9.6* | 42 | -1 | 53 | 9.3* |
| **S8** | 36 | -76 | 2 | 15.3* | 51 | -1 | 50 | 8* |
| **S9** | 42 | -58 | 5 | 18.3* | 42 | -1 | 47 | 10.3* |
| **S10** | 48 | -76 | 2 | 10.7* | 48 | 2 | 47 | 6.4* |
| **S11** | 42 | -64 | 2 | 16* | 51 | -1 | 47 | 12* |
| **S12** | 45 | -64 | 11 | 12.5* | 45 | -4 | 44 | 10.2* |
| **S13** | 45 | -67 | -1 | 30.4* | 39 | -4 | 50 | 17* |
| **S14** | 45 | -61 | -1 | 17.3* | 48 | 5 | 44 | 8.7* |
| **S15** | 48 | -67 | 5 | 12.8* | 45 | 2 | 47 | 10.3* |
| **S16** | 48 | -73 | -1 | 13.3* | 39 | -1 | 50 | 10.3* |
| **S17** | 45 | -64 | -4 | 14.3* | 45 | 5 | 50 | 15* |
| **S18** | 48 | -67 | 2 | 12.9* | _42_ | _-1_ | _47_ | _11.5*_ |
| **S19** | _48_ | _-64_ | _8_ | _17.3*_ | _48_ | _-1_ | _41_ | _11.5*_ |
| **MEAN** | **44.8** | **-67** | **1.5** | **13.5** | **46.6** | **0.7** | **46.2** | **10.4** |
| **± SD** | 3.4 | 5.8 | 4 | 5.5 | 5.2 | 3.9 | 6.2 | 2.5 |

ramps), continuous pursuit with blanking (TRIBL, 18.7°/s, ±15° amplitude, blanked 300 to 1000 ms after ramp onset, four blocks of each seven leftwards and rightwards ramps, preceded by one continuous triangular wave), foveopetal step-ramps (SR, eight blocks with step-ramps directed either leftwards or rightwards per block, eight ramps per block, 18.7 °/s, ± 15° amplitude, ± 2.5° step size, inter-trial central fixation jittered between 1 and 1.5 s), and oscillating pursuit with a stationary random dot background (four blocks of 40 s; red dot with 0.5° radius oscillating at 0.2 Hz, ± 15° amplitude; background: 70 stationary white dots with size 0.5°radius and 2.5° spacing in between background dots) and central fixation with random dots oscillating horizontally in the background (four blocks of 40 s; central fixation red dot, 0.5° radius; background: 70 white dots with radius 0.5° moving at 0.2 Hz and with 2.5° spacing in between dots).

Resulting cartesian MNI-coordinates for the target location estimates are shown in Tab. S1. By default, BOLD activity during continuous pursuit informed the individual definition of V5 and FEF. However, in a few cases (V5: N = 2; FEF: N = 5) no individual activation during continuous pursuit was observed relative to central fixation. In these cases, the combined contrast between all blocks that included SP and the preceding fixation intervals was computed to define V5 and FEF in the right hemisphere. Like the individual V5 locations, the local maxima of estimated individual FEF locations showed considerable variability of 8 ± 5 mm Euclidean distance to the average MNI coordinate at 47/1/46 ± 5/4/6 (M ± SD).

Individual target orientations were estimated using beamforming for the gray matter source space grid points closest to the functional MRI target locations in the V5 and FEF of the right hemisphere following the procedure described in [19]. Across all trials, MEEG datapoints in the time window between 0.3 to 1.3 s relative to ramp onset were extracted and concatenated. Covariance matrices ($\boldsymbol{C}_i$ with $i$ = 1, …, 1000) were computed for each of 1000 bootstrap samples of these MEEG datapoints. A linearly constrained minimum variance (LCMV) beamformer with Unit-Noise-Gain constraint (regularization parameter $\lambda$ = 5%) [19] was used to estimate the orientation of V5 and FEF as the direction of maximum power [20], respectively. In short, the analysis was performed separately for the EEG and MEG data using the complete lead field for the EEG, but only the quasi-tangential components of the lead field for the MEG, reduced by singular value decomposition [21], to regularize the weak radial MEG source orientation components [22]. We then recombined the quasi-tangential MEG source orientations with the radial EEG source orientation, to obtain a combined MEEG source orientation estimate in each bootstrap sample. Across the 1000 bootstrap samples, the medians for each direction (x, y, z) were extracted, and the median vector was normalized to the vector length to estimate the target orientation in V5 and FEF, respectively.
Custom MATLAB scripts were employed using the PsychToolbox for stimulus presentation [23,24], SPM12 [3] for MRI data preprocessing and analysis, and FieldTrip for MEEG data preprocessing and analysis [25].

**S4 Computation of personalized tDCS montages**

The distributed constrained maximum intensity (DCMI) algorithm [26–29] was employed to compute the personalized tDCS montage that maximizes the injected current at the target location along the target orientation (i.e., the directionality). Specifically, the current density vector field $\boldsymbol{j}$ at each node $\boldsymbol{r}_i$ ($i$ = 1, …, $n$) of the FEM head model for a given stimulation montage is determined by the product of the geometry matrix $\boldsymbol{A}$ (dimensions: 3$n$ x ($m$-1)) and the injected current vector $\boldsymbol{s}$ (($m$-1) x 1 vector) applied to $m$-1 non-reference electrodes. $\boldsymbol{A}$ is a matrix where $\boldsymbol{a}_k(\boldsymbol{r}_i)$ contains the 3 x 1 current density vector at node $i$ given a unit strength (1 mA) stimulation from the $k$-th stimulation electrode ($k$ = 1, …, $m$-1) versus the reference electrode (-1 mA) at electrode $m$:

$$\boldsymbol{j} = \boldsymbol{A}\boldsymbol{s} \tag{1}$$

with $\boldsymbol{j} = \begin{bmatrix} \boldsymbol{j}(\boldsymbol{r}_1) \\ \boldsymbol{j}(\boldsymbol{r}_2) \\ \vdots \\ j(\boldsymbol{r}_n) \end{bmatrix}$, $\boldsymbol{A} = \begin{bmatrix} \boldsymbol{a}_1(\boldsymbol{r}_1) & \boldsymbol{a}_2(\boldsymbol{r}_1) & \dots & \boldsymbol{a}_{m-1}(\boldsymbol{r}_1) \\ \boldsymbol{a}_1(\boldsymbol{r}_2) & \boldsymbol{a}_2(\boldsymbol{r}_2) & \dots & \boldsymbol{a}_{m-1}(\boldsymbol{r}_2) \\ \vdots & \vdots & \ddots & \vdots \\ \boldsymbol{a}_1(\boldsymbol{r}_n) & \boldsymbol{a}_2(\boldsymbol{r}_n) & \dots & \boldsymbol{a}_{m-1}(\boldsymbol{r}_n) \end{bmatrix}$ and $\boldsymbol{s} = \begin{bmatrix} \boldsymbol{s}_1 \\ \boldsymbol{s}_2 \\ \vdots \\ \boldsymbol{s}_{m-1} \end{bmatrix}$

[3] https://www.fil.ion.ucl.ac.uk/spm

This formulation can be used to define the optimal injection currents at the stimulation electrodes $\boldsymbol{s}_{max}$: Considering $\boldsymbol{C}$ as the 3 x ($m$-1) submatrix of $\boldsymbol{A}$ that reflects the mapping of the electrode currents to the current density at the target vector, the DCMI is defined as:

$$\boldsymbol{s}_{max} = \arg\max_{\boldsymbol{s}} < \boldsymbol{Cs}, \boldsymbol{o}_t > -\lambda \|\boldsymbol{s}\|_2 \tag{2}$$

subject to $\|\boldsymbol{s}\|_1 \leq 2i_{Total}$ and $\|\boldsymbol{s}\|_\infty \leq i_{Limit}$

with $\boldsymbol{C} = [\boldsymbol{a}_1(\boldsymbol{r}_t), \boldsymbol{a}_2(\boldsymbol{r}_t), \ldots, \boldsymbol{a}_{m-1}(\boldsymbol{r}_t)]$

where $r_t$ denotes target location and $o_t$ the target orientation. We set the overall applied current to $i_{Total}$ = 2 mA (safety constraint) and limited the current per electrode to $i_{Limit}$ = 1.5 mA to reduce transcutaneous tDCS side-effects. The regularization term $\lambda$ serves to minimize the electric field intensity in non-target brain regions and penalizes large values of $\boldsymbol{s}$, effectively spreading the injected current over as many electrodes as possible while maintaining a high directionality within the target. This leads to a well-posed problem with a unique solution which smoothly depends on the initial conditions [26]. $\lambda$ was determined individually for each subject: $\boldsymbol{s}_{max}$ was computed with increasing $\lambda$ (starting with $\lambda$ = 0) until $\boldsymbol{s}_{max}$ contained a total of eight non-zero entries, i.e., eight tDCS electrodes, resulting in $\lambda$ = 0.012 ± 0.018. In addition to the maximum number of eight tDCS electrodes, few occurrences of currents smaller than 0.1 mA for one electrode were distributed among the other same polarity electrodes, due to technical limitations of the tDCS device. Resulting montages with injection currents $\boldsymbol{s}_{max}$ described personalized cathodal tDCS montages.

To quantify individual transcranial electric field parameters, for each subject, we computed the target intensity ($|E|_{IT} = \|\sigma^{-1}\boldsymbol{Cs}_{max}\|_2$), the target intensity corrected for the parallelity between the stimulation target orientation vector and the target electric field orientation vector (directionality, $|E|_{DIR} = < \sigma^{-1}\boldsymbol{Cs}_{max}, \boldsymbol{o}_t >$) [27–29], as well as the spatial extent of the electric field relative to the stimulation target ($|E|_{EXTENT}$) [28,29]. The overall cortical electric field was computed as $|\mathbf{E}(\boldsymbol{r}_i)| = |\sigma^{-1}\boldsymbol{j}_{max}(\boldsymbol{r}_i)|$, given $\boldsymbol{j}_{max} = \boldsymbol{As}_{max}$.

### S5 Personalized tDCS targeting the right FEF

In experiment 2, personalized tDCS was applied targeting the right FEF (Fig. S1). Individual FEF estimates were determined by functional MRI data (location) and combined MEEG (orientation) during continuous SP. Individual tDCS montages were computed and applied as described in the main manuscript. Average electrode montages for the estimated FEF targets showed a cluster of anodes in right frontal, frontocentral, central and parietocentral electrodes, and a frontal/frontocentral, as well as a parietal cathodal cluster. Average electric fields are well targeted to the FEF in the right hemisphere.

### S6 Normal electric field of personalized tDCS targeting the right V5

To illustrate the applied electric field for personalized cathodal tDCS targeting the right V5, individual normal electric fields were computed. For the individual FEM head models, the normal to the cortical surface was computed as the unit length vector between each gray

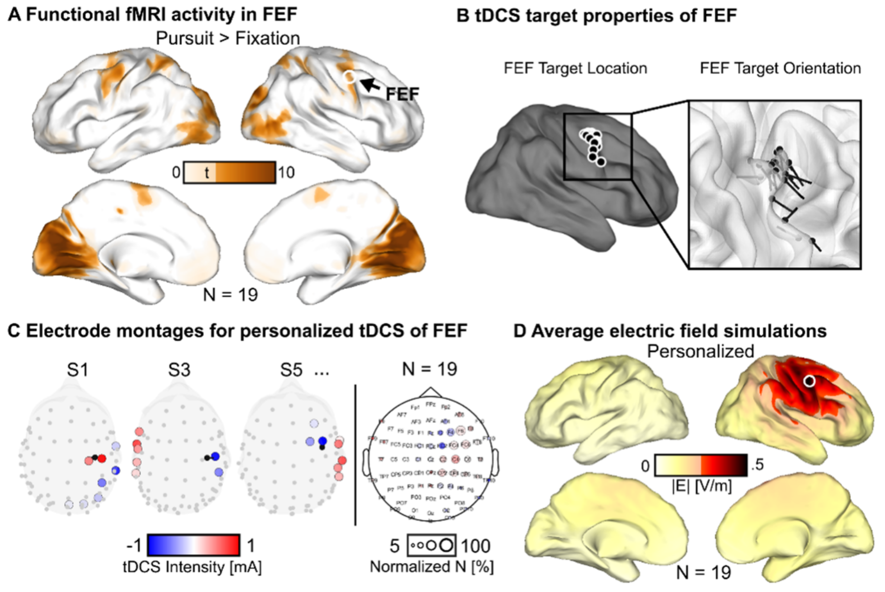


**Figure S1. Personalized tDCS targeting FEF. A)** Second-level functional MRI BOLD activity during ongoing SP, compared to fixation, revealed a significant cluster that corresponds to the area FEF. Significant *t*-values are shown (FDR-corrected), interpolated on the inflated cortical surface of the MNI brain. **B)** Individual tDCS target area FEF. Locations (left) are shown on an inflated cortical surface of the MNI brain. Details of FEF orientations (right) are shown together with the white matter surface of the MNI brain. **C)** Personalized tDCS montages targeting individual FEF are shown for three exemplary subjects (left; top view) and for the whole sample in a topographical representation (right). **D)** Lateral and medial view of average simulated electric fields targeting the right FEF interpolated on the inflated cortical surface of the MNI brain, with values below 50% of the maximal intensity masked (light colors). The average tDCS target location in the right FEF is indicated by a white circle.

matter node coordinate and the closest white matter node coordinate. The normal electric fields $\mathbf{E}_{normal}$ were computed as the dot product between each normal vector with the respective electric field vector (Fig. S2). Overall, normal electric fields indicate average inward currents in right posterior cortex but show high inter-individual variability and limited contribution to the overall electric field. This might be explained by the individual orientations of target V5 that also show a high variability following the cortical structure of V5. As depicted in the main manuscript (Fig. 2B), only in a few cases these estimated summed dipole orientations are quasi-radial but rather they are quasi-tangential or in between a clear radial or tangential orientation.

### S7 Eye tracking data acquisition and analysis for the tDCS experiments

A video-based eye tracker was employed to record eye movements (Eyelink 1000Plus, SR Research Ltd., Ottawa, Canada; Eyelink host software version 5.17; heuristic single-stage filter [30]) in a closed room with lights off. Participants were placed 65 cm in front of an LCD monitor (XL2720, BenQ, Taipeh, Taiwan; 1920 x 1080 pixel, i.e., 49.3° x 28.9° visual size; 120 Hz refresh rate) on a chin-forehead rest. To ensure a calibration error smaller than 1° visual angle, binocular 13-point calibration (calibration point positions [x, y] in pixel: [960, 540], [384, 216], [960, 108], [1536, 216], [576, 324], [1344, 324], [192, 540], [1728, 540], [576, 756], [1344, 756], [384, 864], [960, 972], [1536, 864]) was performed and validated. SP performance was assessed before, during and after 20 minutes of tDCS ($t_0$, $t_{TDCS\_5}$, $t_{TDCS\_10}$, $t_{TDCS\_15}$, $t_{TDCS\_20}$, $t_{15}$, $t_{40}$), as described in the main manuscript. During $t_{TDCS}$, SP task blocks were separated by simple oculomotor tasks to activate the oculomotor system during the stimulation, while providing an active rest for the participants at the same time. Simple oculomotor tasks consisted of short video clips [31], as well as continuous oscillating pursuit with stationary background (60 s duration; red dot of size 0.5° oscillating at 0.2 Hz, ± 15° amplitude; background: 70 stationary white dots with size 0.5° and 2.5° spacing) and fixation with moving background (60 s duration; central fixation of a red dot, size 0.5°; background: 70 white dots with size 0.5° (2.5° spacing) moving at 0.2 Hz). Stimuli were presented by custom MATLAB software (R2019b, The Mathworks Ltd., Natick, MA, USA) using the PsychToolbox (version 3.0.16) [23,24].

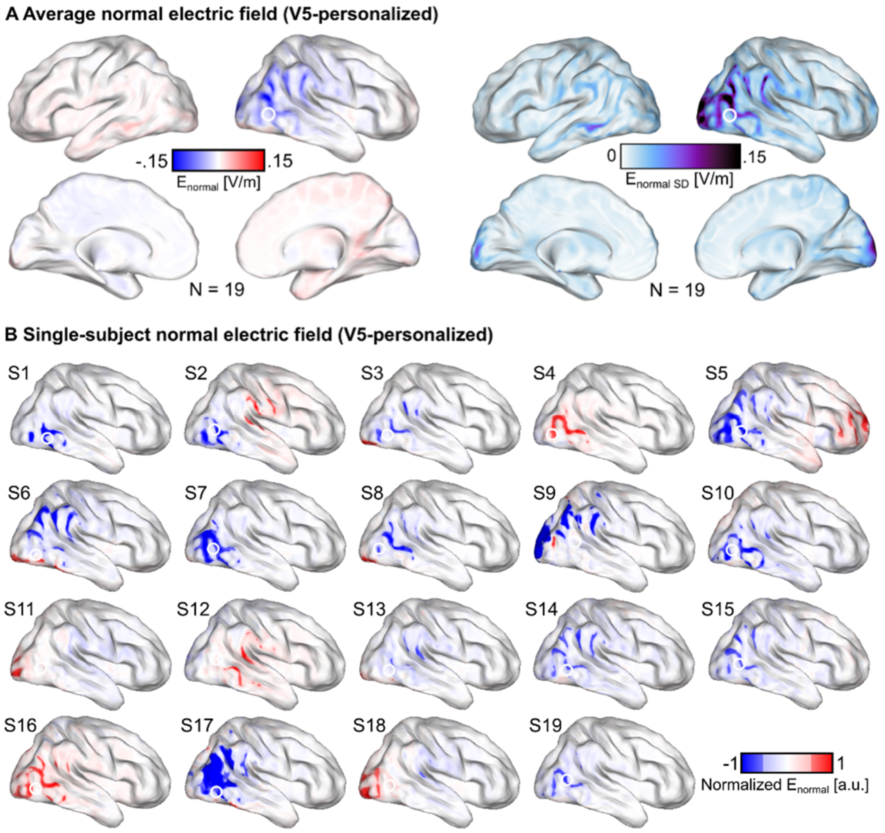


**Figure S2. Normal electric fields for personalized tDCS targeting V5. A)** For personalized tDCS targeting V5 the mean (left) and standard deviation (right) of personalized transcranial normal electric fields $\mathbf{E}_{normal}$ targeting V5 are shown on the inflated cortical surface of the MNI brain. **B)** Single-subject personalized normal electric fields $\mathbf{E}_{normal}$ are shown on the inflated cortical surface of the MNI brain. White circles illustrate the average V5 location.

A custom semi-automatic pre-processing procedure [32] was applied to transfer raw gaze coordinate data of one eye (by default the left eye) to visual angle data. The right eye was only used in rare cases of limited data quality restricted to the left eye (experiment 1 (V5): 2.5 ± 5.8 %, experiment 2 (FEF): 1 ± 3.4 % of SP task blocks). Continuous data were lowpass-filtered at 50 Hz using a Gaussian filter and SP velocities were computed as the first derivative of eye position (average of ± 8 ms before and after any given data point). Data intervals holding saccades (experiment 1 (V5): 2.04 ± 0.56 saccades/s, experiment 2 (FEF): 2.09 ± 0.51 saccades/s), blinks (experiment 1 (V5): 0.08 ± 0.07 blinks/s, experiment 2 (FEF): 0.08 ± 0.04 blinks/s), and invalid data were removed. Continuous data were epoched (-100 to 1600 ms for TRI and TRIBL relative to the left or right ramp onset and -100 to 1300 ms for SR relative to the ramp onset) and invalid epochs were rejected (experiment 1 (V5): 1.5 ± 2.1 %, experiment 2 (FEF): 0.5 ± 0.6 % of epochs were rejected). Median eye velocity traces across epochs were computed separately for each subject, tDCS condition (anodal, cathodal, sham), timepoint ($t_0$, $t_{TDCS\_5}$, $t_{TDCS\_10}$, $t_{TDCS\_15}$, $t_{TDCS\_20}$, $t_{15}$, $t_{40}$) and stimulus direction (leftward, rightward). The correct automatic detection of eyeblinks, saccades and intervals of artefactual signal was validated manually. Data quality was good (experiment 1 (V5): 1.04 ± 0.8 % missing data; 284 ± 54 %

signal-to-noise ratio; experiment 2 (FEF): 1.18 ± 0.6 % missing data; 297 ± 87 % signal-to-noise ratio).

SP parameters were computed for open-loop SP initiation during the SR task (see main manuscript) and closed-loop SP maintenance (Fig. S3). Open-loop SP initiation was quantified by computing the SP latency, and initial acceleration from median eye velocities obtained during the SR task. Initial acceleration was determined as the slope of a regression line fitted to the eye velocity trace after the movement onset of the visual target stimulus. Specifically, multiple robust regressions were fitted to different time windows of the eye velocity trace starting at the timepoint where the velocity exceeded 3.2 SDs of baseline eye velocity for subsequent 20 ms [33]. The model best fitting the initial velocity changes was selected, and the initial acceleration was defined based on the slope of this regression line. SP latency was defined as the time of the intercept between the same regression line with the baseline velocity plateau before the movement onset of the target visual stimulus [32].

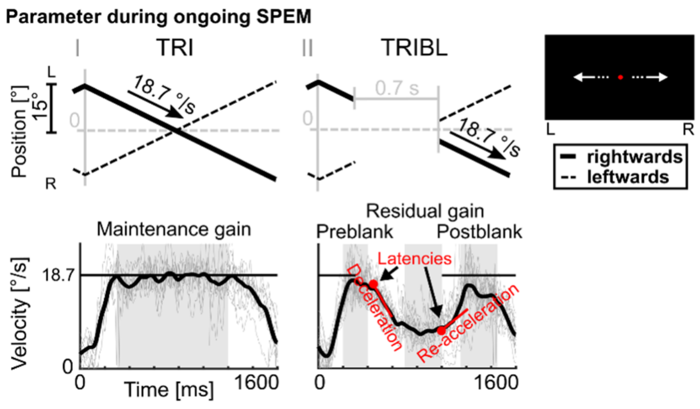


**Figure S3. SP parameters during ongoing pursuit.** During tDCS experiments, three tasks were conducted to assess different aspects of smooth pursuit eye movements (SP). Besides foveopetal step-ramps (SR; see main manuscript), continuous pursuit (TRI) and continuous pursuit with blanking (TRIBL) were conducted. Top: Horizontal eye position as a function of time is shown for leftward and rightward ramps. Roman numerals indicate epoch intervals from figure 1 in the main manuscript. Bottom: Single-trial (thin gray lines) and median velocity traces (black lines) computed across single-trials are depicted for one exemplary subject the TRI and TRIBL tasks. Intervals for the velocity gain computation are indicated by gray shaded areas. For TRI, maintenance gain was computed to quantify the predictive SP performance with ongoing visual input (300 to 1200 ms). In TRIBL trials, velocity gain was computed before target disappearance (preblank gain, 200 to 400 ms), during blanking (residual gain, 700 to 1000 ms), and after reappearance of the visual target (postblank gain, 1150 to 1450 ms). During TRIBL, deceleration following blanking onset, and re-acceleration before reappearance are indicated by red regression lines (deceleration and re-acceleration are estimated as slopes of these regression lines). The respective latencies (i.e., deceleration latency and re-acceleration latency) are indicated by red dots. Target velocity is indicated by a black line at 18.7 °/s.

Closed-loop SP maintenance during ongoing movement of a visual target stimulus (TRI) was defined by computing the maintenance gain as the ratio of the median eye velocity across samples between 300 to 1200 ms (relative to ramp onset) and target velocity. For SR, the early maintenance gain was computed (300 to 700 ms relative to target movement onset). Similarly, preblank velocity gain (200 to 400 ms relative to ramp onset), residual velocity gain (700 to 1000 ms) and postblank velocity gain (1150 to 1450 ms) were computed as estimates of the SP maintenance before, during and after target blanking during the TRIBL task. Furthermore, for TRIBL eye deceleration following target disappearance and re-acceleration after target re-appearance were determined as the slope of a regression line fitted to eye velocity trace after target disappearance (deceleration) and target re-appearance at the end of the blanking interval (re-acceleration), respectively. Deceleration and re-acceleration latencies were defined as the time of the intercept between the same regression lines with the preblank and residual eye velocity plateaus after the disappearance (deceleration latency) or before the re-appearance (re-acceleration latency) of the blanked visual target.

#### S8 Baseline pursuit velocities before tDCS application

Baseline SP velocities before tDCS application at $t_0$ are shown to illustrate the overall sanity of eye movement recordings and good performance across participants. For visualization, SP median velocity traces were lowpass-filtered at 10 Hz and mean and 95%-confidence intervals across subjects were computed (Fig. S4).

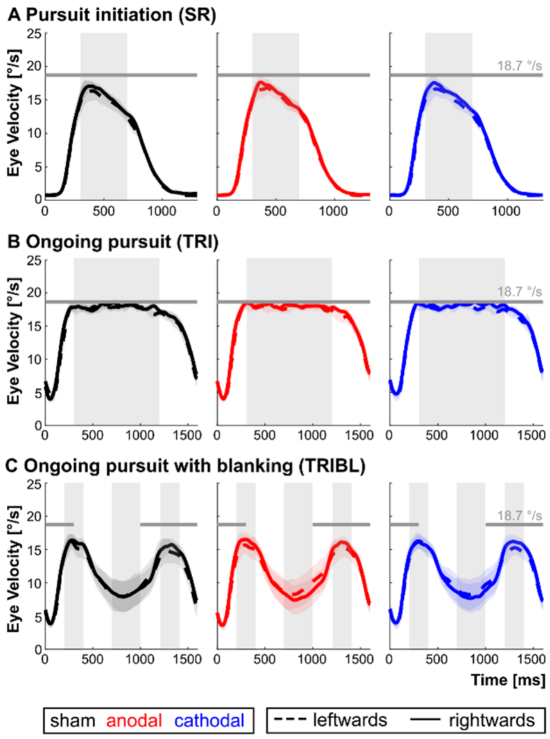


**Figure S4. Baseline SP velocities before tDCS application.** Grand average and SEM across single-subject median velocity traces before application of sham, anodal, and cathodal tDCS ($t_0$) and for leftwards and rightwards SP, respectively. Velocity traces are shown for **A)** SP initiation and early closed-loop SP during the SR task, **B)** ongoing SP during the TRI task (middle), and **C)** SP during the TRIBL task (bottom). Target velocity is depicted by a gray line at 18.7 °/s. Gray shaded areas indicate latency intervals for the computation of early maintenance gain (A), maintenance gain (B), as well as preblank, residual, and postblank gain (C).

#### S9 The influence of tDCS side-effects

During the application of tES, side-effects can occur that may alter or even compromise the transcranial effect of tES [compare 34]. To evaluate the influence of tDCS side-effects, subjective transcutaneous side-effects (after ramp-up of tDCS) were rated by participants after each tDCS session. In addition, the subjective fatigue was retrospectively rated after each tDCS session for four timepoints during the tDCS, corresponding to the SP measurements at $t_{TDCS\_5}$, $t_{TDCS\_10}$, $t_{TDCS\_15}$, and $t_{TDCS\_20}$. Overall low side-effects were reported for somatosensory (M ± SD = 0 ± 0 to 0.1 ± 0.37, i.e. absent) and pain perception (M ± SD = 0 ± 0 to 0.2 ± 0.54, i.e. absent; Fig. S5). Between tDCS conditions no significant difference was observed for the subjective somatosensory ($X^2{}_2$ = 5.6, $p$ = .111) or pain perception ($X^2{}_2$ = 4.7, $p$ = .222). However, the Friedman ANOVA revealed significant fatigue differences between measurement timepoints ($X^2{}_2$ = 73.8, $p$ < .001). As indicated by post-hoc Wilcoxon tests, no differences between tDCS conditions were observed (all $|Z|$ ≤ 1.7, all $p$ ≥ .148). However, participants significantly reported increased fatigue during the second half of $t_{TDCS}$ (M ± SD = 2.3 ± 0.93 to 2.8 ± 1.03, i.e. awake to moderate fatigue), compared to the initial wakefulness in all three tDCS conditions (M ± SD = 1.5 ± 0.51 to 1.7 ± 0.67, i.e. awake; all $|Z| \geq 2.9$, all $p \leq .035$). In addition, during the anodal session similarly increased fatigue was reported during $t_{TDCS\_15}$, compared to the second quarter $t_{TDCS\_10}$ ($Z$ = -3.6, all $p$ = .002).

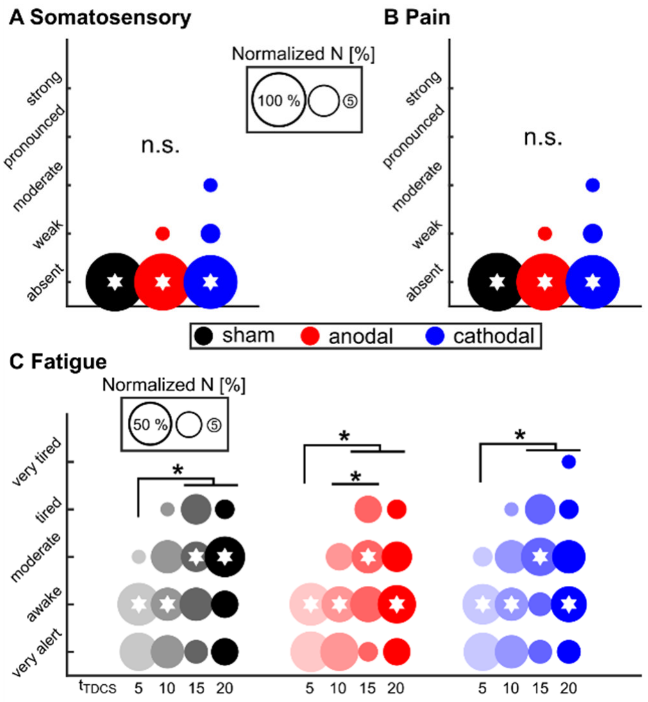


**Figure S5. Side-effects of tDCS. A)** Subjective ratings of somatosensory side-effects during tDCS are depicted for sham, anodal, and cathodal tDCS. Only few participants reported weak to moderate somatosensory side-effects during anodal and cathodal tDCS. No significant difference was observed between tDCS conditions. **B)** Subjective ratings of pain side-effects during tDCS are depicted for sham, anodal, and cathodal tDCS. Only few participants reported weak to moderate pain side-effects during anodal and cathodal tDCS. No significant difference was observed between tDCS conditions. **C)** Fatigue was subjectively rated for four timepoints during $t_{TDCS}$. No significant difference was observed between tDCS conditions. However, participants described significantly increased fatigue for the second half of the 20 minutes tDCS application, compared to the initial timepoint in all tDCS conditions. In addition, participants described increased fatigue during $t_{TDCS\_15}$, compared to $t_{TDCS\_10}$ during anodal tDCS. * indicate $p < .05$ (Bonferroni-corrected).

Based on these results, the main effect of covariate fatigue and the interaction between fatigue and the timepoint during $t_{TDCS}$ were included as fixed effects in the LMM analysis of SR latency to validate the observed results. However, no main ($F_{1,232}$ = 1.6, p = .205) or interaction effect of fatigue was revealed ($F_{3,261}$ = 0.7, p = .569) and improvements by adding fatigue as covariate in the statistical $model_A$ ($AIC_A$ = 3591), compared to the basic $model_0$ without including side-effects ($AIC_0$ = 3605) were negligible. Thus, no side-effects were considered in the final statistical analysis of SP latencies in the SR task.

### S10 Individual electric field simulations

Individual electric field simulations were computed and interpolated on the cortical surface of the MNI brain for personalized tDCS targeting V5 (Fig. S6) and normative tDCS over V5 (Fig. S7).

### S11 Practice-effects and fatigue-effects during experiment 1 (V5-personalized)

During typical tDCS paradigms, tDCS is applied for several minutes and the effect of tDCS is read-out during and after tDCS and compared to the behavioral performance or neurophysiological activity before the application of tDCS ($t_0$). However, due to the same experimental design changes over time may occur irrespective of modulations due to tDCS. In the present experiment 1 (V5-personalized) SP was significantly modulated as a function of timepoint for several SP parameters (Tab. S3), in addition to the observed tDCS online effect on SP latencies that is indicated by the interaction between tDCS condition (sham, anodal,

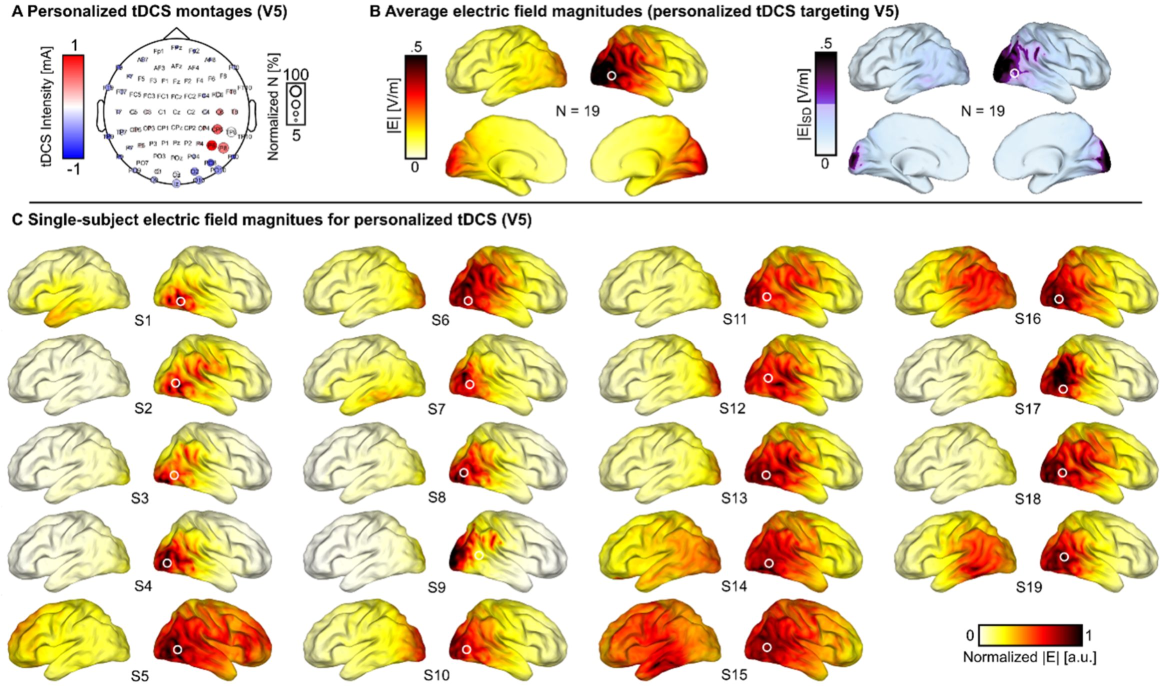


**Figure S6. Single-subject electric field simulations for personalized tDCS targeting V5. A)** For personalized tDCS targeting V5 the average montage topography is shown including sensor labels. Color indicates the average intensity across all N = 19 subjects for each sensor. Sensor sizes indicate how often the respective electrode was part of the personalized montage across subjects. **B)** Mean (left) and standard deviation (right) of personalized transcranial electric field magnitudes |**E**| targeting V5 are shown on the inflated cortical surface of the MNI brain. **C)** Single-subject personalized electric field magnitudes |**E**| are shown on the inflated cortical surface of the MNI brain.

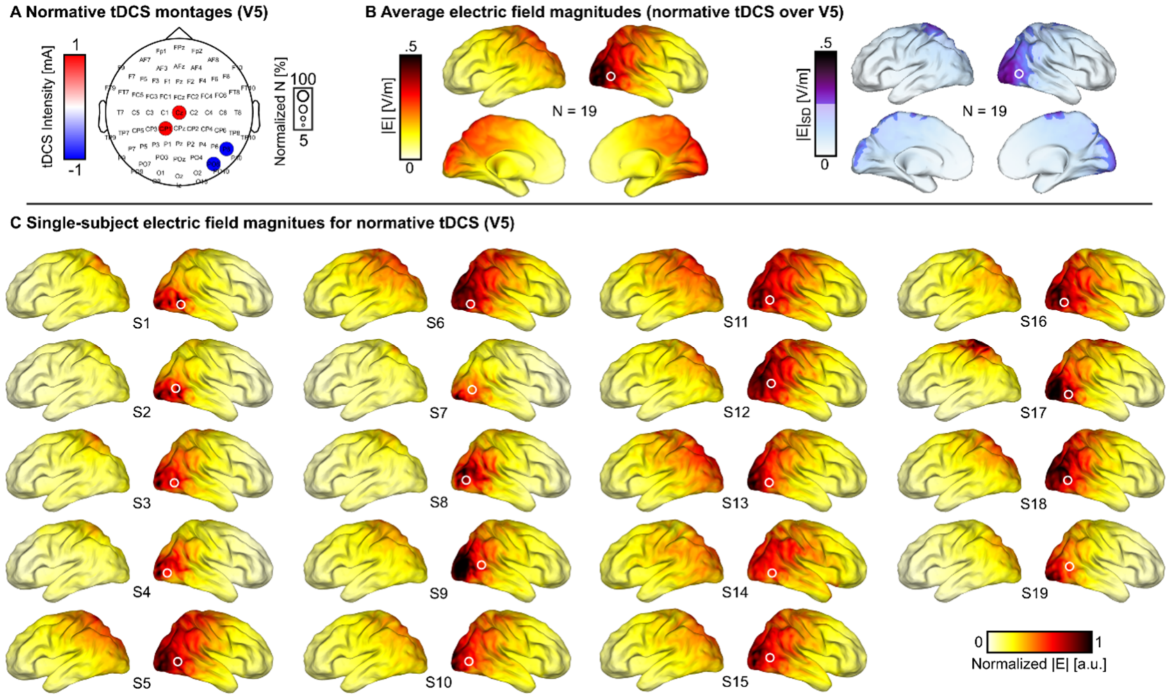


**Figure S7. Single-subject electric field simulations for normative tDCS over V5. A)** For normative tDCS over V5 the average montage topography is shown including sensor labels. Color indicates the average intensity across all N = 19 subjects for each sensor. Sensor sizes indicate how often the respective electrode was part of the normative montage across subjects (i.e., the same montage was applied across subjects). **B)** Mean (left) and standard deviation (right) of normative transcranial electric field magnitudes |**E**| over V5 are shown on the inflated cortical surface of the MNI brain. **C)** Single-subject normative electric field magnitudes |**E**| are shown on the inflated cortical surface of the MNI brain.

cathodal), timepoint ($t_0$, $t_{TDCS\_5}$, $t_{TDCS\_10}$, $t_{TDCS\_15}$, $t_{TDCS\_20}$, $t_{15}$, and $t_{40}$) and stimulus direction (leftwards, rightwards).

Specifically, in line with a previous study [1], timepoint main effects were observed for the maintenance gain during ongoing SP (TRI), the residual gain, re-acceleration latency, and postblank gain during ongoing SP with target blanking (TRIBL), as well as for initial acceleration and early maintenance gain during the SR task. In contrast to a previous report [1], we observed a timepoint main effect for the preblank gain, but no timepoint main effect for deceleration and re-acceleration during the blanking task (TRIBL). We add to previous findings, that in the present experiment 1 (V5-personalized), SP was analyzed at four timepoints during tDCS application ($t_{TDCS\_5}$, $t_{TDCS\_10}$, $t_{TDCS\_15}$, $t_{TDCS\_20}$), in addition to before and after tDCS ($t_0$, $t_{15}$, $t_{40}$). Specifically, we report a significant improvement of SP performance after tDCS ($t_{40}$), compared to preceding SP before and during tDCS ($t_0$, all $t_{TDCS}$ timepoints) for the maintenance gain (all $p \leq .001$), the residual gain (all $p \leq .003$), the postblank gain (all $p \leq .002$; also $t_{15} > t_0$: $p < .001$), and for the early maintenance gain (all $p \leq .001$). Preblank gain was less clearly increased during both $t_{15}$ and $t_{40}$ only compared to later $t_{TDCS}$ timpoints ($t_{TDCS\_15}$, $t_{TDCS\_20}$; all $p \leq .012$). Re-acceleration was improved for $t_{40}$ compared to $t_0$, $t_{TDCS\_10}$, and $t_{TDCS\_15}$ (all $p \leq .027$). Initial acceleration was improved for $t_{40}$ compared to all $t_{TDCS}$ timpoints (all $p \leq .01$). However, during tDCS performance was significantly decreased, compared to $t_0$ before and/or $t_{15}$ after tDCS for maintenance gain ($t_{TDCS\_15} < t_0$: $p = .022$; $t_{TDCS\_10}$ to $t_{TDCS\_20} < t_{15}$: all $p \leq .028$), for preblank gain ($t_{TDCS\_15}$ and $t_{TDCS\_20} < t_{15}$: all $p \leq .012$), residual gain ($t_{TDCS\_10}$ to $t_{TDCS\_20} < t_{15}$: all $p \leq .001$), postblank gain ($t_{TDCS\_10}$ to $t_{TDCS\_20} < t_{15}$: all $p \leq .001$), initial acceleration ($t_{TDCS\_20} < t_0$: $p = .005$; $t_{TDCS\_20} < t_{15}$: $p = .003$), and for early maintenance gain ($t_{TDCS\_10}$ to $t_{TDCS\_20} < t_0$: all $p \leq .017$; $t_{TDCS\_5}$ to $t_{TDCS\_20} < t_{15}$: all $p \leq .003$). Taking together, we observed both SP practice-effects that improved performance within experimental sessions, but also SP fatigue-effects during the period of 20 minutes tDCS.

Within-session practice-effects have been reported previously and were suggested to rely on top-down extraretinal SP mechanisms [35–40] due to the clear improvement of SP in the absence of visual input during blanking [1]. These findings are in line with the present improvements during the TRIBL task. The same top-down mechanisms might also affect be involved in the improvements observed during SP with continuously visible target during the TRI task [1] and do not contradict otherwise reported stable SP performance over larger time scales that provide evidence for SP as a stable biomarker in healthy participants and patients suffering from psychosis spectrum disorder [41–49].

Considering the relatively long period of 20 minutes tDCS, it is reasonable to assume that fatigue affected the SP performance during $t_{TDCS}$. The analysis of subjective ratings provide evidence for the increasing fatigue during tDCS, especially during the latter 10 minutes of stimulation (see S9). It must be noted that previous findings described the critical role of functionally specific endogenous brain activity during direct current stimulation to produce Hebbian learning-like neuronal modulation [50]. In other words, tDCS might modulate those functions only or more effectively that show endogenous activation. Following this assumption, we designed the experiment including SP tasks during tDCS application to induce endogenous activity the same SP brain network, including V5, that we wanted to target with personalized tDCS. However, performing SP over more than 5 minutes, even with active rest periods (simple oculomotor tasks and video clips), increased subjective fatigue ratings, especially for the latter 10 minutes of tDCS. Therefore, we conclude that the observed modulation of ipsiversive SP latencies by personalized cathodal tDCS are not the product of the transcranial electric field per se, but rather the interplay between tDCS and the endogenous brain activity during $t_{TDCS}$ that itself is dynamically shaped by practice and fatigue. Beside the control of the transcranial electric fields by personalized applications, the understanding of endogenous brain activity during tDCS remains an important factor to control tDCS outcomes.

**S12 Differences between pursuit stimulus directions in experiment 1 (V5-personalized)**

Independent of tDCS modulations as described in the main manuscript, effects of SP target direction were observed across various parameters during all three tasks (TRI, TRIBL, SR). Direction main effects were observed for the maintenance gain during ongoing SP (TRI), the preblank gain, deceleration latency, deceleration and postblank gain during ongoing SP with target blanking (TRIBL), as well as for SP latencies and early maintenance gain during the SR task (Tab. S3). For most of these parameters, we observed a significantly better performance for rightwards SP, compared to leftwards SP (maintenance gain, preblank gain, deceleration, postblank gain, early maintenance gain: all R > L, all $p < .001$; SP latency: R < L, $p < .001$). In line with previous reports [51–53], functional MRI of the participants in this study indicated more pronounced activity in V5 of the right hemisphere (see main manuscript). When foveating a single SP target stimulus as in the present SP tasks, increased V5 activity in the MRI might indicate an increased foveal representation of direction-sensitive neurons in MT that have been shown to facilitate ipsiversive SP when modulated with microstimulation [54] and produce impaired ipsiversive SP when lesioned [55].

Only for deceleration latency (R < L, $p = .019$) and re-acceleration latency (R > L, $p = .019$) during the TRIBL task an inverted direction effect was observed. These differences mark an earlier decay of prediction at the beginning of the blanking interval and later anticipation before the re-occurrence of the target in rightwards, compared to leftwards SP [1,35,36]. These findings are partly in line with previous reports of earlier and facilitated re-acceleration during SP with blanking [1]. The authors suggested an interpretation of the better performance for leftwards SP in these isolated parameters as an indicator of asymmetries in visuo-spatial attention that is biased to the left hemifield due to a right hemispheric lateralization of the underlying neural network [56,57]. During SP, it has been shown that visuo-spatial attention constantly shifts ahead of the foveated visual target [58]. Thus, the lateralization of visuo-spatial to the right hemisphere might explain why leftwards SP might benefit from visuo-spatial attention more than rightwards pursuit in a retinotopic matter [59,60]. Visuo-spatial attention might especially gain importance for the otherwise decreased performance of leftwards, compared to rightwards SP.

### S13 Experimenter instructions and subjective tDCS perception across experiments

Participants in all three experiments were blind to the applied tDCS procedure and received only general information in written form prior to participation that allowed informed consent and the adherence to safety and exclusion criteria for the applied methods. Identical task instructions were presented in printed form prior to each experimental task and for all participants after initial setup of eytracking, tDCS, MRI, and/or MEEG to reduce a potential experimenter bias to the minimum.

In addition, we quantified the subjectively rated transcutaneous perception after each tDCS application (compare S9) and compared these ratings across tDCS conditions (sham, anodal, cathodal) and experiments (1: V5-personalized, 2: FEF-personalized, 3: V5-normative). These ratings were used to control differences across tDCS conditions that might bias participants performance by providing structured perceptual clues about the tDCS conditions or montages. In all three experiments, the average subjective ratings of tDCS perception were negligible (experiment 1, V5-personalized: somatosensory M ± SD = 0 ± 0 to 0.1 ± 0.37, pain M ± SD = 0 ± 0 to 0.2 ± 0.54, i.e. absent; experiment 2, FEF-personalized: somatosensory M ± SD = 0 ± 0 to 0.1 ± 0.27, pain M ± SD = 0 ± 0 to 0.1 ± 0.23, i.e. absent; experiment 3, V5-normative: somatosensory M ± SD = 0.1 ± 0.19 to 0.2 ± 0.3, pain M ± SD = 0.1 ± 0.32 to 0.2 ± 0.54, i.e. absent). Importantly, neither experiment showed distinct ratings with respect to somatosensory or pain perception across tDCS conditions as assessed by separate Friedman ANOVAs (all ${X^2}_2 \leq 5.6$, $p \geq .111$). Also, statistical analysis did not reveal a difference of subjective tDCS perception between experiment 1 and 2 (i.e., V5-personalized compared to FEF-personalized; Friedman ANOVAs, all ${X^2}_5 \leq 8.8$, $p \geq .114$), and no difference between experiment 1 and 3 (i.e., V5-personalized compared to V5-normative; Bonferroni-corrected Mann-Whitney-U tests, all $|Z|_{36} \leq 1.9$, $p \geq .573$). We conclude that participants were not able to dissociate tDCS conditions and specifics of the applied tDCS montage based on perceptual cues introduced by the tDCS procedure. Taking together the results from all three experiments, the observed results can be considered specific with respect to pursuit initiation as a subfunction of smooth pursuit, to the targeted V5 brain region, as well as to the application of personalized tDCS targeting V5.

**Table S2. Personalized tDCS targeting V5 modulates pursuit initiation.** Results of LMM analysis indicate significant modulation of SP initiation by personalized tDCS targeting V5 (considering timepoints $t_0$, $t_{TDCS\_5}$, $t_{TDCS\_10}$, $t_{TDCS\_15}$, $t_{TDCS\_20}$, $t_{15}$, and $t_{40}$). TRI = continuous pursuit, TRIBL = continuous pursuit with blanking, SR = step-ramp. * indicate *p* < .05.

| Task | Parameter | tDCS condition | | tDCS condition * Timepoint | | tDCS condition * Direction | | tDCS condition * Timepoint * Direction | |
|---|---|---|---|---|---|---|---|---|---|
| | | F | *p* | F | *p* | F | *p* | F | *p* |
| **TRI** | Maintenance gain | 0.1 | *.99* | 0.84 | *.605* | 0.975 | *.378* | 0.549 | *.882* |
| **TRIBL** | Preblank gain | 1.21 | *.301* | 1.68 | *.068* | 0.87 | *.42* | 0.43 | *.953* |
| | Deceleration latency | 1.67 | *.191* | 0.4 | *.964* | 0.36 | *.696* | 0.67 | *.778* |
| | Deceleration | 1.14 | *.32* | 0.38 | *.971* | 0.23 | *.797* | 0.66 | *.79* |
| | Residual gain | 0.1 | *.902* | 0.34 | *.981* | 0.38 | *.684* | 0.15 | *> .999* |
| | Re-acceleration latency | **8.69 *** | ***< .001*** | 1.11 | *.348* | 0.36 | *.702* | 0.83 | *.617* |
| | Re-acceleration | 0.17 | *.845* | 0.22 | *.998* | 0.17 | *.843* | 0.92 | *.523* |
| | Postblank gain | 1.21 | *.299* | 0.86 | *.591* | 0.04 | *.958* | 0.31 | *.988* |
| **SR** | Pursuit latency | 0.64 | *.53* | 0.78 | *.675* | 0.37 | *.693* | **1.81 *** | ***.044*** |
| | Initial acceleration | 0.11 | *.9* | 0.59 | *.848* | 0.08 | *.921* | 1.61 | *.085* |
| | Early maintenance gain | 2.78 | *.064* | 0.55 | *.882* | 0.35 | *.706* | 0.42 | *.956* |

**Table S3. Main effects of timepoint and pursuit stimulus direction.** Results of linear mixed model analysis for each estimated oculomotor parameter during experiment 1 (V5-personalized), considering timepoints $t_0$, $t_{TDCS\_5}$, $t_{TDCS\_10}$, $t_{TDCS\_15}$, $t_{TDCS\_20}$, $t_{15}$, and $t_{40}$. TRI = continuous pursuit, TRIBL = continuous pursuit with blanking, SR = step-ramp. * indicate *p* < .05.

| Task | Parameter | Timepoint | | Direction | | Timepoint * Direction | |
|---|---|---|---|---|---|---|---|
| | | F | *p* | F | *p* | F | *p* |
| **TRI** | Maintenance gain | **12.73 *** | ***< .001*** | **43.28 *** | ***< .001*** | 0.804 | *.567* |
| **TRIBL** | Preblank gain | **5.47 *** | ***< .001*** | **17.17 *** | ***< .001*** | 0.51 | *.802* |
| | Deceleration latency | 1.71 | *.118* | **5.55 *** | ***.019*** | 0.57 | *.752* |
| | Deceleration | 1.02 | *.412* | **12.64 *** | ***< .001*** | 1.06 | *.387* |
| | Residual gain | **15.76 *** | ***< .001*** | 3.73 | *.055* | 0.34 | *.916* |
| | Re-acceleration latency | **5.03 *** | ***< .001*** | **11.08 *** | ***.001*** | 1.2 | *.307* |
| | Re-acceleration | 0.78 | *.583* | 0.16 | *.688* | 1.67 | *.128* |
| | Postblank gain | **18.34 *** | ***< .001*** | **50.83 *** | ***< .001*** | 0.25 | *.959* |
| **SR** | Pursuit latency | **13.42 *** | ***< .001*** | **31.02 *** | ***< .001*** | 0.68 | *.664* |
| | Initial acceleration | **8.9 *** | ***< .001*** | 0.4 | *.527* | 0.64 | *.698* |
| | Early maintenance gain | **26.03 *** | ***< .001*** | **31.41 *** | ***< .001*** | 0.82 | *.556* |

**Table S4. No online effects by personalized tDCS targeting the right FEF.** Results of LMM analysis do not indicate a significant modulation of SP parameters by personalized tDCS targeting FEF during the tDCS application (online effects, i.e. considering timepoints $t_{TDCS\_5}$, $t_{TDCS\_10}$, $t_{TDCS\_15}$, and $t_{TDCS\_20}$). TRI = continuous pursuit, TRIBL = continuous pursuit with blanking, SR = step-ramp. * indicate *p* < .05.

| Task | Parameter | tDCS condition | | tDCS condition * Timepoint | | tDCS condition * Direction | | tDCS condition * Timepoint * Direction | |
|---|---|---|---|---|---|---|---|---|---|
| | | F | *p* | F | *p* | F | *p* | F | *p* |
| **TRI** | Maintenance gain | 2.09 | *.128* | 0.94 | *.467* | 0.09 | *.915* | 0.21 | *.975* |
| **TRIBL** | Preblank gain | 2.2 | *.113* | 0.33 | *.92* | 0.24 | *.787* | 0.32 | *.926* |
| | Deceleration latency | 1.38 | *.256* | 0.96 | *.454* | 1.56 | *.213* | 0.98 | *.438* |
| | Deceleration | 1.05 | *.353* | 0.38 | *.892* | 0.22 | *.803* | 0.36 | *.903* |
| | Residual gain | **7.72 *** | ***.001*** | 0.62 | *.716* | 0.28 | *.758* | 0.21 | *.974* |
| | Re-acceleration latency | 0.51 | *.602* | 0.43 | *.86* | 0.3 | *.742* | 0.47 | *.83* |
| | Re-acceleration | 1.11 | *.331* | 0.56 | *.759* | 2 | *.137* | 1.12 | *.35* |
| | Postblank gain | **8.7 *** | **< .001** | 0.63 | *.71* | 0.62 | *.541* | 0.16 | *.986* |
| **SR** | Pursuit latency | 0.31 | *.737* | 0.73 | *.628* | 0.25 | *.78* | 0.43 | *.862* |
| | Acceleration | 1.71 | *.184* | 1.76 | *.106* | 1.49 | *.228* | 0.48 | *.824* |
| | Early maintenance gain | 1.36 | *.259* | 0.61 | *.727* | 2.57 | *.079* | 0.37 | *.9* |

**Table S5. No online effects by normative tDCS over V5.** Results of LMM analysis do not indicate a significant modulation of SP parameters after normative tDCS over V5 (online effects, i.e. considering timepoints $t_{TDCS\_5}$, $t_{TDCS\_10}$, $t_{TDCS\_15}$, and $t_{TDCS\_20}$). TRI = continuous pursuit, TRIBL = continuous pursuit with blanking, SR = step-ramp. * indicate *p* < .05.

| Task | Parameter | tDCS condition | | tDCS condition * Timepoint | | tDCS condition * Direction | | tDCS condition * Timepoint * Direction | |
|---|---|---|---|---|---|---|---|---|---|
| | | F | *p* | F | *p* | F | *p* | F | *p* |
| **TRI** | Maintenance gain | 0.66 | *.521* | 0.91 | *.486* | 0.03 | *.971* | 0.2 | *.976* |
| **TRIBL** | Preblank gain | **4.02 *** | ***.02*** | 0.54 | *.775* | 0.23 | *.799* | 0.63 | *.708* |
| | Deceleration latency | 0.21 | *.811* | 0.87 | *.518* | 1.12 | *.33* | 1.29 | *.26* |
| | Deceleration | 0.2 | *.822* | 0.46 | *.841* | 0.3 | *.743* | 0.49 | *.818* |
| | Residual gain | **5.27 *** | ***.006*** | 0.47 | *.833* | 0.24 | *.787* | 0.44 | *.854* |
| | Re-acceleration latency | 0.22 | *.804* | 1.38 | *.223* | 0.24 | *.785* | 0.96 | *.452* |
| | Re-acceleration | 3 | *.053* | 0.93 | *.475* | 1.19 | *.307* | 0.54 | *.776* |
| | Postblank gain | 2.28 | *.106* | 0.3 | *.938* | 2.6 | *.077* | 0.33 | *.922* |
| **SR** | Pursuit latency | 2.26 | *.108* | 0.21 | *.973* | 1.31 | *.273* | 0.31 | *.93* |
| | Acceleration | 0.06 | *.994* | 0.23 | *.968* | 0.21 | *.813* | 0.76 | *.6* |
| | Early maintenance gain | 1.11 | *.332* | 1.14 | *.555* | 0.04 | *.965* | 0.23 | *.967* |
